# Long-term patient-derived ovarian cancer organoids closely recapitulate tumor of origin and clinical response

**DOI:** 10.1101/2025.01.10.631512

**Authors:** L. Thorel, E. Dolivet, P-M. Morice, R. Florent, J. Divoux, M. Perréard, L. Lecouflet, G. Desmartin, C. Marde Alagama, F. Giffard, A. Leconte, J. Lequesne, B. Clarisse, M. Briand, A. Traore, C. Villenet, JP. Meneboo, G. Babin, L. Gaichies, S. Martin-Françoise, JF. Le Brun, R. Rouzier, E. Brotin, C. Denoyelle, N. Vigneron, R. Leman, D. Vaur, L. Castera, C. Blanc-Fournier, N. Elie, B. Plancoulaine, F. Joly, M. Meryet-Figuière, M. Figeac, L-B. Weiswald, L. Poulain

**Author notes:** Co-corresponding authors: Louis-Bastien Weiswald, INSERM U1086 ANTICIPE (Interdisciplinary Research Unit for Cancers Prevention and Treatment), Comprehensive Cancer Center François Baclesse, 3 Avenue du Général Harris, BP 45026, 14 076, Caen cedex 05, France. Laurent Poulain, INSERM U1086 ANTICIPE (Interdisciplinary Research Unit for Cancers Prevention and Treatment), Comprehensive Cancer Center François Baclesse, 3 Avenue du Général Harris, BP 45026, 14 076, Caen cedex 05, France. These authors equally contributed to this work (L-B. Weiswald, L. Poulain).

## Abstract

There is an urgent need of precision medicine for ovarian cancer patients to identify patients who respond to chemotherapy and PARP inhibitors, a therapy targeting homologous recombination deficiency (HRD). Here we established a panel of 37 long-term patient-derived tumor organoids (PDTO) models of various histological subtypes from 224 patients and demonstrated that they mimic the histological and molecular characteristics of original tumors. Screening of chemotherapeutic drugs showed that PDTO exhibit heterogeneous responses, and that response of PDTO from high-grade serous ovarian carcinoma to carboplatin recapitulated patient response to first-line treatment. Additionally, the detection of HRD phenotype of PDTO using functional assay was associated with the results of the HRD test Genomic Instability Scar (GIScar). Although larger-scale investigations are needed to confirm the predictive potential of PDTO, these results provide further evidence of the potential interest of ovarian PDTO for functional precision medicine even if many challenges remain to be addressed.

## Introduction

Ovarian cancer (OC) is the 8th most common cancer in women worldwide, with approximately 325,000 new cases and 207,000 deaths in 2022^1^. Epithelial ovarian cancers represent 95% of ovarian cancers and are a heterogeneous disease. Based on histopathology and molecular genetic alterations, they are divided into 5 major subtypes with distinct biological and molecular properties: high-grade serous, low-grade serous, clear cell, endometrioid, and mucinous ovarian carcinomas^2^. Due to the usually asymptomatic nature of early stages, most of OC cases are diagnosed at late stages (eg. stage III/IV as determined by the International Federation of Obstetrics and Gynecology (FIGO)) when cancer has spread widely to the peritoneal cavity^3^. The standard of care consists of primary or interval cytoreductive surgery followed by combination platinum-taxane chemotherapy with or without bevacizumab^4^. However, around 70% of patients with OC will recur after first-line therapy, leading to a 5-year relative survival rate of only 29% for late stages^5^. Thus, it is well known that a “one-size-fits-all” treatment approach should no longer be applied. The introduction of targeted maintenance therapy with poly ADP-ribose polymerase inhibitors (PARPi) to the first-line setting has led to remarkable improvement in outcome in selected patients, especially those with tumors harboring *BRCA1/2 (breast cancer genes 1 and 2)* mutations and homologous recombination deficiency (HRD)^6^. At time of recurrence, according to the guidelines, the choice of treatment regimen will be based on platinum sensitivity, prior lines of treatment, chemotherapy-related adverse events, *BRCA* mutation status and physician and patients’ preferences^4^. In the case of early recurrence, the options of treatment are limited, with a response rate of less than 15%^2^, highlighting the lack of predictive biomarkers for guided anticancer agent selection. This also applies to PARPi, since the sensitivity to PARPi is not restricted to tumors harboring *BRCA1/BRCA2* mutations^7^. Therefore, a range of genetic tests are being developed to improve profiling of HR status, including the academic HRD test Genomic Instability Scar (GIScar)^8^.

However, genomics precision medicine has several limitations, such as the lack of selectivity of some molecular signatures^7^, the limits of the interpretation (complex mutational signatures or variants of unknown signification)^9^ and the fact that genomic instability signatures inform on a history of instability. Moreover, it does not warrant that such instability is still present.

This underscores the interest in developing functional precision medicine, an approach based on direct drug exposure of patient-derived tumor models to predict patient response^12^. Improved feasibility of generating preclinical models mimicking the patient tumor, such as patient-derived tumor organoids (PDTO), has made these models accessible for personalized treatment. PDTO are 3D multicellular structures generated from patient tumor cells embedded in basement membrane matrix and cultured in a medium supplemented with a cocktail of cell signaling pathways activators and inhibitors to reproduce *in vivo* niche conditions and allow long term growth. PDTO faithfully recapitulate the histological and molecular characteristics of the tumor from which they are derived. They can be rapidly expanded and established from a small sample size such as needle biopsy with higher success rate compared to other models^13^. Most of all, a growing body of evidence indicates that PDTOs are able to predict the clinical response, although most of clinical studies were based on small sample size (<10 patients)^14,15^. PDTO cultures have already been successfully established from ovarian cancer patients^16–24^ with success rate of establishment ranging from 18% to 90% but with varied definitions of an established model. However, in most of the studies, the success rate of establishment as well as long-term stable passage is usually not explicitly mentioned in the literature^25^. Interestingly, several of them demonstrated high level of correlation between response of PDTO and clinical response despite the small sizes of samples, ranging from 1 to 7 patients^20,22–24^.

In this study, we established and performed a comprehensive characterization of a panel of 37 long-term PDTO derived from 31 patients with various OC subtypes and determined whether response of PDTO to first-line chemotherapy correlates with patient response to assess clinical relevance of these models.

## Results

### Establishment and characterization of long-term ovarian PDTO

Long-term PDTO were considered established as PDTO line after cell expansion over 8 successive passages and the ability to grow in culture after thawing (13.8% success rate, n=224, Supplementary Table 1). We generated 37 ovarian cancer long-term PDTO from 31 patients of various histotypes, with a majority of high-grade serous ovarian cancer (HGSOC, 23/31) but also rarer subtypes, including ovarian clear cell carcinomas (OCCC, 4/31), carcinosarcoma (CS, 2/31), endometrioid (EM, 1/31) and mucinous (MC, 1/31) (Figure 1A).

**Figure 1.**
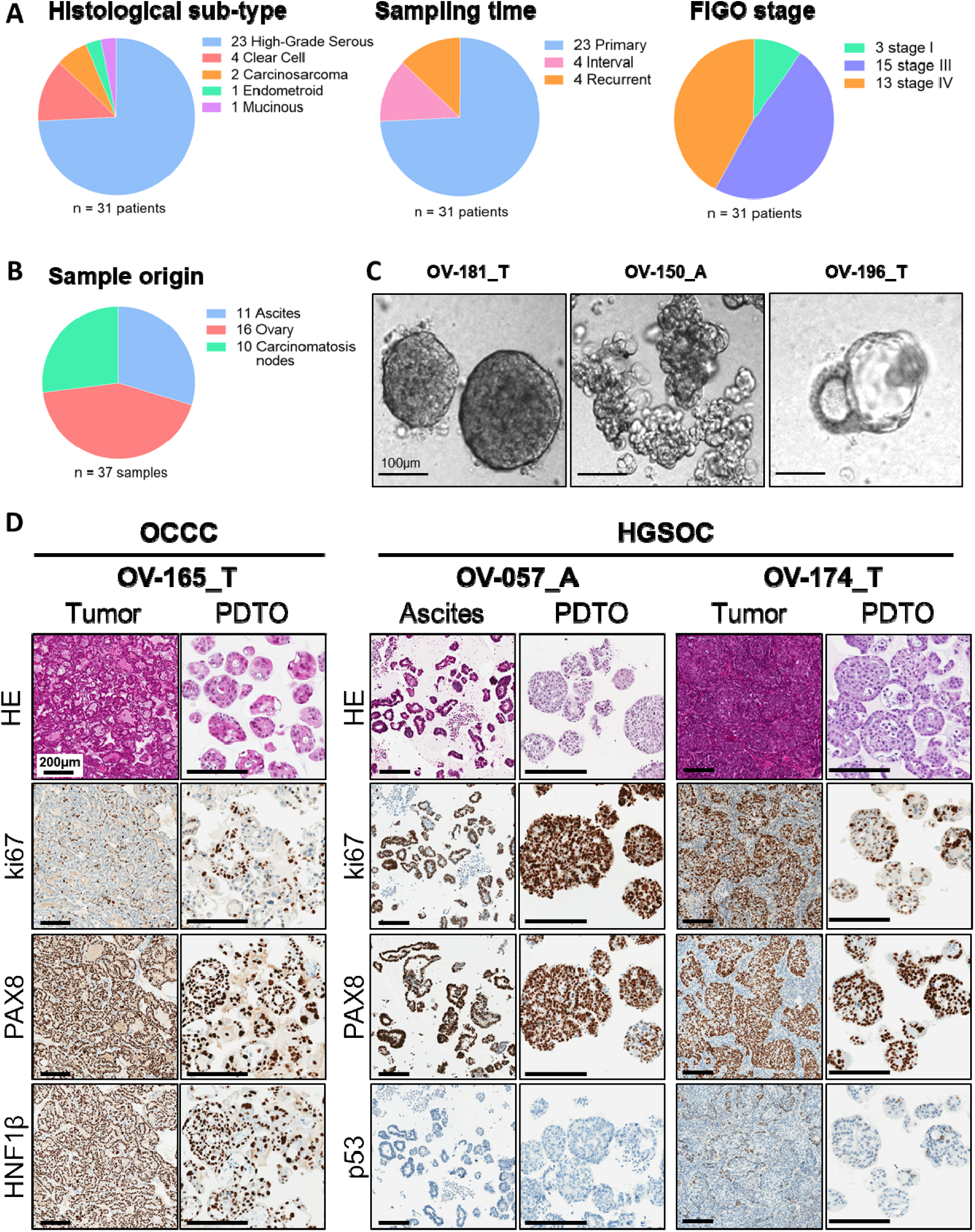
Establishment and characterization of a PDTO panel from ovarian tumors. A. Pie-chart showing the histological subtype, sampling time, sample origin and FIGO stage of the 31 patients from which PDTO lines were established. B. Pie-chart showing the sample origin and of the 37 ovarian PDTO lines. C. Representative pictures of PDTO of various morphologies. A = Ascites-derived PDTO and T = Tumor-derived PDTO. Scale bar = 100pm. D. Hematoxylin and eosin (HE) staining and Immunohistochemistry analysis of ovarian cancer protein markers ki67, PAX8, p53 and HNF1β in tumor tissues and paired PDTO. Scale bar = 200μm.

Out of 37, 8 PDTO were established from pretreated patients. All of these 8 PDTO models were generated from 8 independent patients with HGSOC or CS including 4 from an interval surgery (after 3 chemotherapy cycles) and 4 from ascites received after 2 or more treatment lines (Figure 1A). Moreover, since ovarian cancer is often diagnosed at a late stage, most of the PDTO lines were derived from stage FIGO III and IV (28/31). Stage FIGO I (3/31) are exclusively derived from mucinous and clear cell carcinomas (Figure 1A, Supplementary Table 1). Samples were mainly obtained from solid tumor (26/37 established models, including 16 ovary and 10 carcinomatosis nodes) but PDTO establishment from ascitic samples was also successful (11/37 established models) (Figure 1B). More patient clinical data are presented in Supplementary Table 2.

Ovarian PDTO models displayed heterogeneous phenotypes, such as dense morphology for the majority of cases (Figure 1C, left panel), grape-like (Figure 1C, middle panel), or cystic morphologies (Figure 1C right panel). In order to compare ovarian PDTO with their tumor of origin, hematoxylin and eosin (HE) staining was performed, as well as immunostaining of ovarian cancer diagnosis proteins including PAX8, p53 and a marker of proliferation Ki67.

For OCCC, additional staining of HNF1β was achieved in order to better characterize this specific subtype (Figure 1D)^26^. Overall, PDTO retained the morphological features of their original tumor, as well as the expression profile of ovarian cancer diagnosis proteins.

### Ovarian PDTO recapitulate tumor’s genomic landscape

For 15 PDTO and their matched tumor obtained between 2018 and 2021, low-pass WGS was performed in order to analyze overall copy number variation (CNV) across the genome. Overall, ovarian PDTO models displayed a good level of similarity with their original tumor (Figure 2A). The similarity level of all paired samples varied between 0.4 and 1 with 60% of the pairs above 0.7 (Supplementary Table 3).

**Figure 2.**
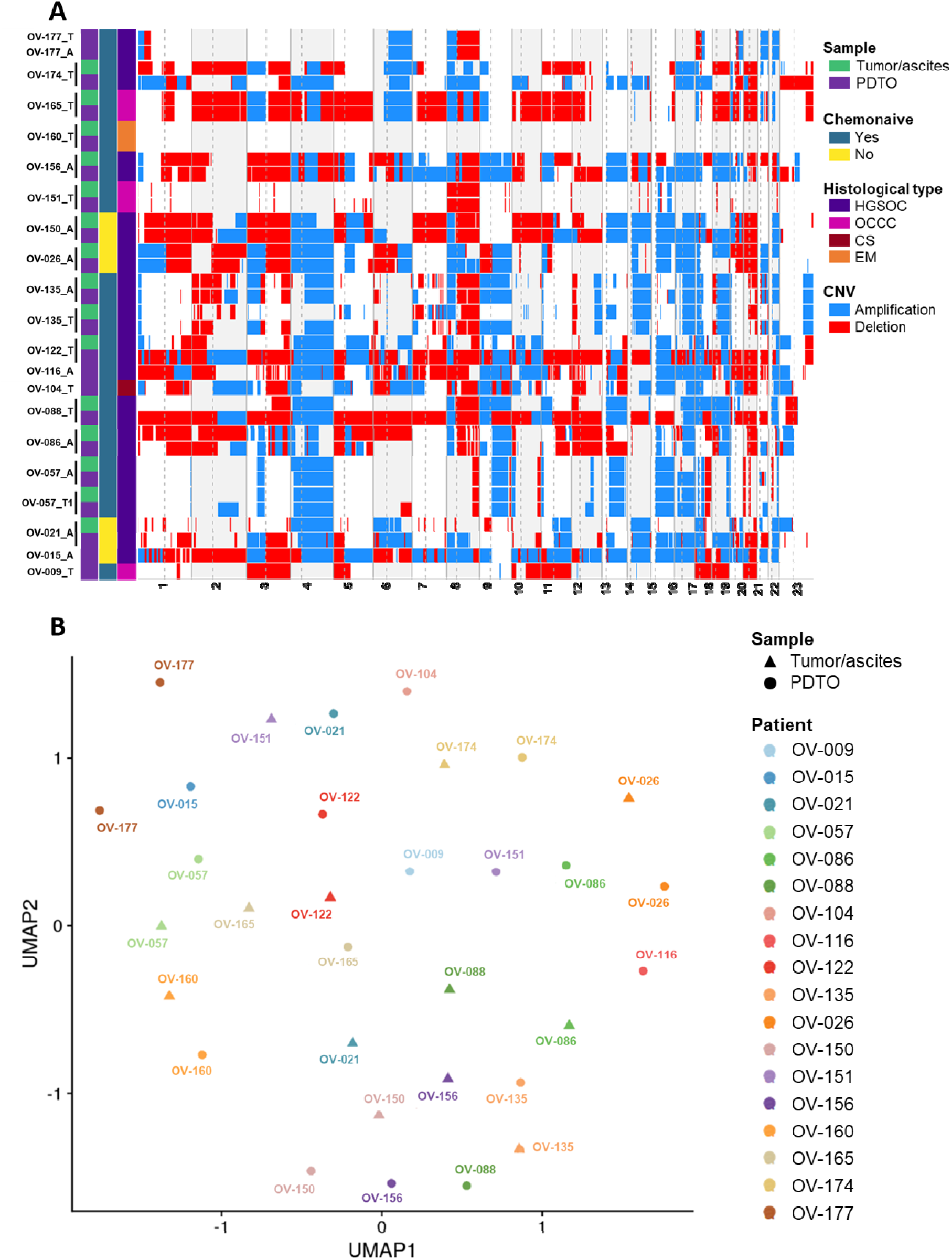
Ovarian PDTO lines captured genomic and transcriptomic features of parental tumor. A. Genome-wide heatmap of DNA copy number gains (blue) and losses (red) of tumor tissues and paired PDTO. A = Ascites-derived PDTO and T = Tumor-derived PDTO. B. Distribution of tumor samples and paired PDTO on UMAP clusters based on gene expression profile following NMF (k=12).

Transcriptomic profiling was performed on the same paired samples (n=15) and displayed a strong correlation between genes expressed by the PDTO and the tumor (R² ranging from 0.613 to 0.827 with a mean of 0.693) (Supplementary Figures 1A and B). However, Uniform Manifold Approximation and Projection (UMAP) visualization of all samples showed a clear split, the tumor samples clustering together away from PDTO samples (Supplementary Figure 1C). Analysis of differentially expressed gene enrichment showed a strong enrichment in immunology-related pathways, consistent with the fact that stromal cell component is lacking in PDTO models (Supplementary Figures 1D and E). In order to better compare paired samples, a non-negative matrix factorization was used, which allowed us to evidence an overlap between tumor and PDTO samples with most of the pairs clustering together (Figure 2B, Supplementary Figures 2A and B). All together these analyses showed that our heterogeneous panel of ovarian PDTO are representative of their tumors of origin.

### PDTO drug response correlation to patient clinical outcome

Next, we performed drug-response profiling on 29 ovarian PDTO for which a clinical follow-up of more than 12 months was available, representing 26 patients out of 31 in order to explore the potential of PDTO to recapitulate patient clinical response to chemotherapy. Since carboplatin is the standard-of-care for all ovarian cancers, all PDTO were exposed to increasing concentrations of this drug and response was assessed using viability assay.

We selected a homogenous sample of subjects among our cohort of patients to provide a meaningful analyse. Therefore, only PDTO derived from chemonaive HGSOC patients were considered. Patients who received immunotherapy during the first line treatment were excluded, since they were part of clinical trials and no double-blind release was done (Figure 3A). The selected PDTO (n=9) displayed heterogenous response to carboplatin with IC50 (half inhibitory concentration) ranging from 5.43 to 51.64 µM and AUC (area under the curve) from 1260 to 5819 (Figures 3B and C). Normalized AUC z-score of the PDTO response to carboplatin was then plotted as a violin diagram (Figure 3D) showing a clear split between sensitive and resistant PDTO around the value 0. When compared to the patient clinical outcome, the PDTO response to carboplatin was associated to the patient platinum-free interval (PFI, time between the last carboplatin cycle and the relapse) (Figure 3E). Some discrepancies were found when compared to the progression-free survival (PFS), to the overall survival (OS) and to the CA-125 normalization (nadir <35U/mL) while no correlation at all were found using KELIM score (Figure 3E). Interestingly the most resistant patient, OV-130_A with a PFI inferior of 1 month (therefore considered as platinum-refractory) was by far the most resistant PDTO to carboplatin (Figures 3C, D and E). However, the difference of carboplatin normalized AUC between platinum-sensitive (PFI ≥ 6 months) and platinum resistant (PFI < 6 months) patients did not reach significance (p-value= 0.38) (Figure 3F), notably due to low statistical power and the influence by the OV-135_T and the OV-301_T PDTO models. Both models displayed sensitive response to carboplatin while the patients were considered as platinum-resistant. Despite the initial good response of these patients during the first three cycles of chemotherapy (Supplementary Figure 3A and B), patients OV-135 and the OV-301 relapsed respectively from a distant mediastinal metastasis and a periaortic lymph node. For the patient OV-135, a PDTO model was also established from a chemonaive ascitic sample (OV-135_A), but it displayed the same sensitivity to carboplatin (Supplementary Figures 3C and D). Overall, the response of chemonaive HGSOC PDTO allowed to distinguishing the patients with resistant PDTO (n=5) with a median PFI of 1.61 months from the patients with sensitive PDTO (n=4) with a median PFI of 6.80 months (p-value = 0.06) (Figure 3G).

**Figure 3.**
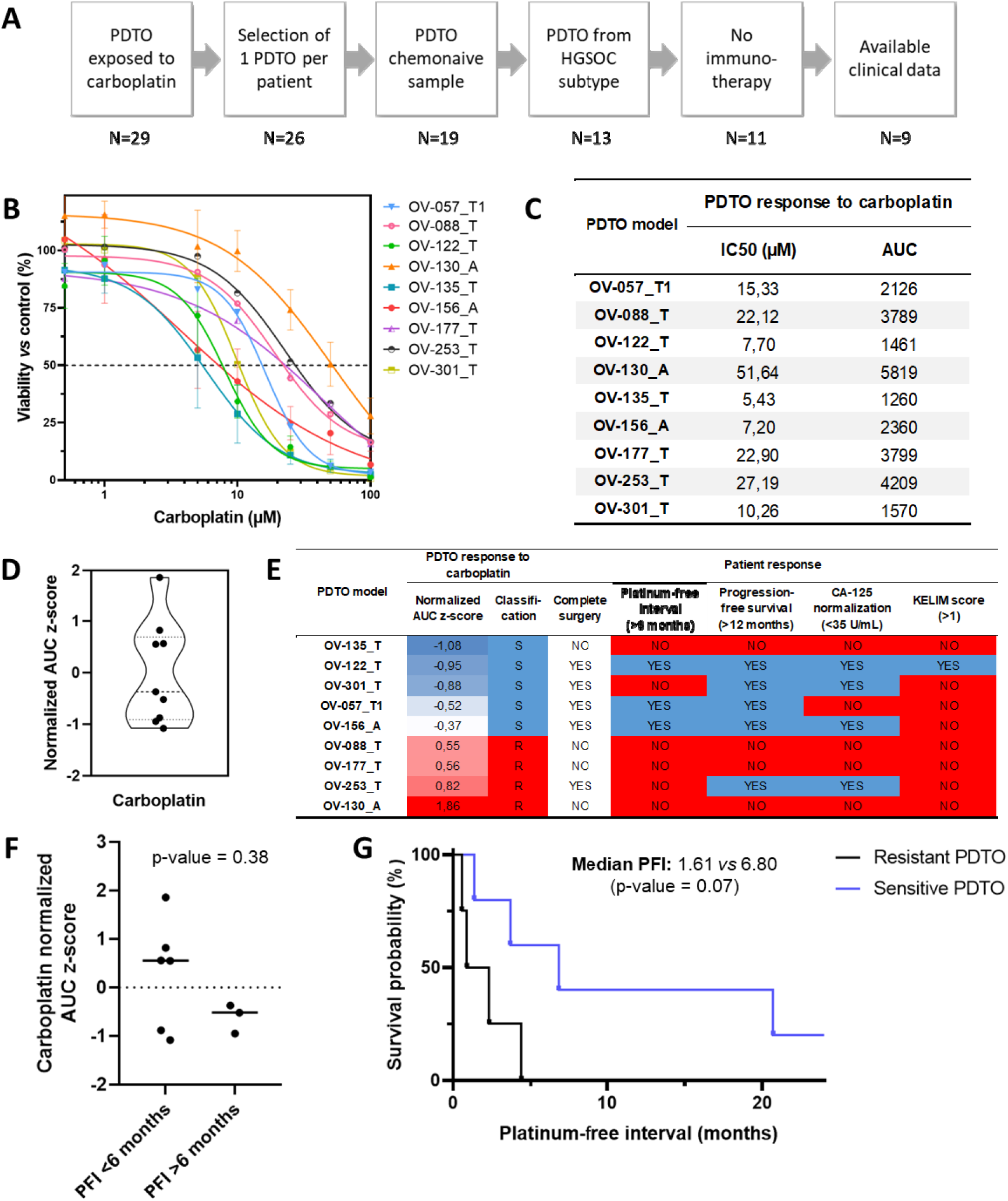
Ovarian PDTO lines recapitulate patient response to carboplatin. A. Flow chart of the PDTO selection for the comparison with patient clinical outcome. B. Dose-response curves of the 9 selected PDTO, each curve is representative of at least two independant experiments. A = Ascites-derived PDTO and T = Tumor-derived PDTO. C. Summary of the IC50 (μM) and AUG obtained In B. for the 9 PDTO models. D. Representation of the carboplatin normalized AUG z-score (n=9). E, Comparison of the PDTO response to carboplatin and clinical outcome, expressed as complete surgery, platinum-free interval (PFI), progression-free survival (PFS), CA-125 normalization and KELIM score. F. Dot plots comparing PDTO response to carboplatin, expressed as normalized AUC z-score, and clinical outcome expressed as PFI. Data were analyzed using unpaired two-sided Mann-Whitney test. G, Kaptan-Meter plot comparing PFI of the resistant PDTO group (n=5) and the sensitive PDTO group (n=4). Data were analyzed using Gehan-Breslow-Wilcoxon test.

Next, our PDTO panel was exposed to a larger number of drugs used in the clinical management of ovarian cancers. To that end, we firstly adapted our protocols in order to work on a smaller number of PDTO per replicate, allowing an easier and faster screening. This optimization was performed from 96-well plate to 384-well plate, and required 4 times less PDTO (50 *vs* 200) (Supplementary Figure 4A). This miniaturization approach was validated on 15 PDTO models and showed a strong correlation between both protocols (Spearman r = 0.83, p = 0.002) (Supplementary Figures 4B and C).

We then tested 6 different drugs, all routinely used in the clinical management of ovarian cancers (1^st^ line, maintenance and following ones), namely, carboplatin, paclitaxel, doxorubicin, gemcitabine and 2 PARP inhibitors (olaparib and niraparib) on our panel of 29 ovarian PDTO (Figure 4A). Globally, the sensitivity to carboplatin was associated to the sensitivity to other drugs, with some exceptions. For example, the PDTO model OV-301_T displayed a sensitivity to all drugs with the exception of the two PARPi. Conversely, the PDTO OV-130_A was resistant to all drugs except to olaparib (Figure 4A).

**Figure 4.**
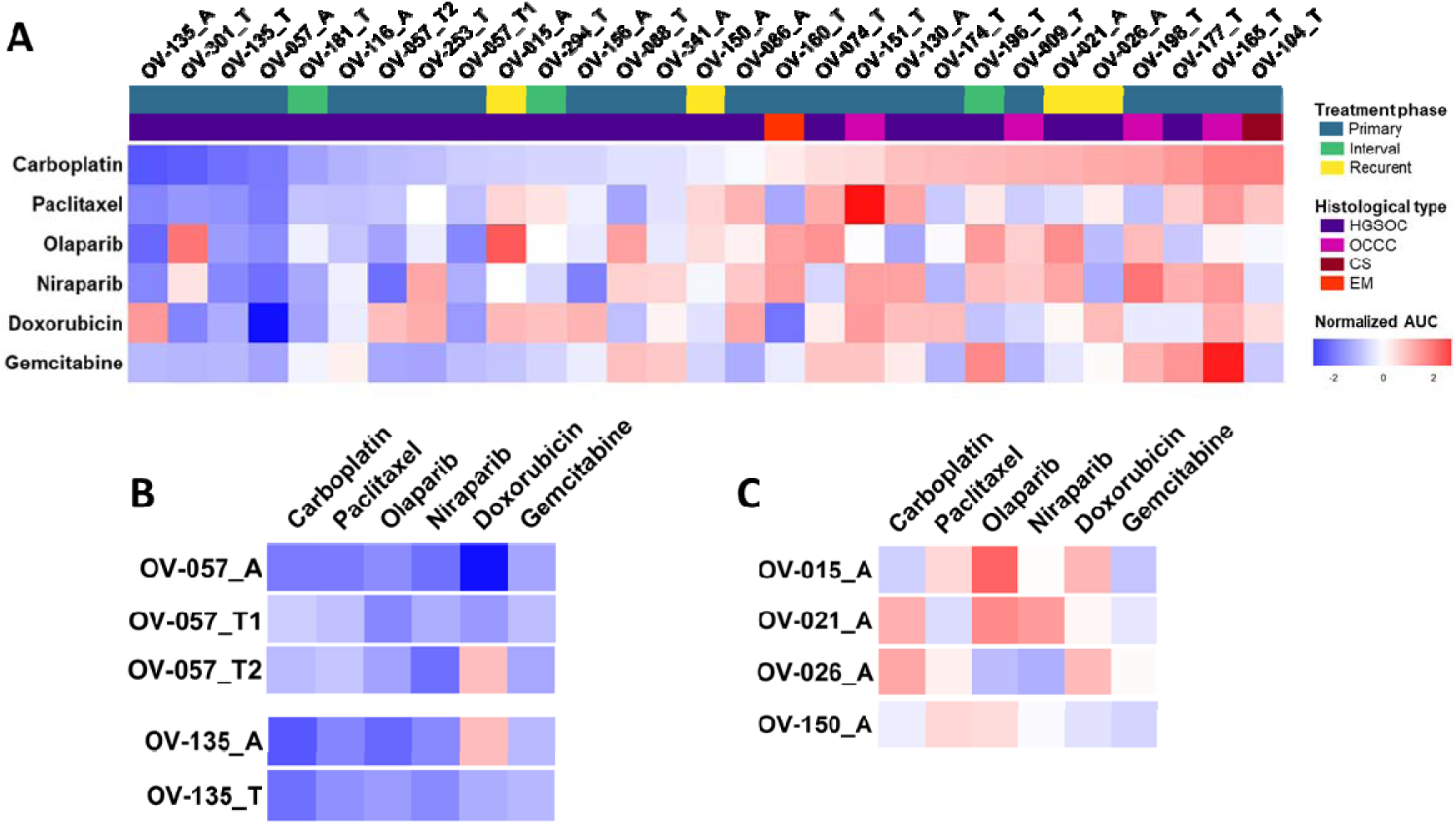
Drug screening of PDTO standard-of-care therapies in ovarian cancer. A. Heatmap showing PDTO response to carboplatin, paclitaxel, olaparib, niraparib, doxorubicin and gemcitabine, expressed as normalized AUC z-score (n=29 models). A = Ascites-de rived PDTO and T = Tumor-derived PDTO. B. Heatmap showing response of PDTO models derived from the same patient (n=2 patients). C. Heatmap showing response of PDTO models derived from recurrent ovarian cancer (n=4 patients).

As expected, all PDTO derived from OCCC and CS were in the platin-resistant side of the heatmap, consistent with the known platin-resistance of these subtypes (Figure 4A).

We were able to derive multiple PDTO models from different tumor and ascitic samples of 2 patients. Regarding the patient OV-057, 3 chemonaives HGSOC PDTO models were established from an ascitic sample (OV-057_A), an epiploon node (OV-057_T1) and from a peritoneal node (OV-057_T2). 2 HGSOC PDTO models were also derived from an ascitic sample and from a peritoneal node of the patient OV-135 (OV-135_T and OV-135_A, respectively). Overall, PDTO models generated from the same patient responded similarly to all the drugs tested, whatever their origin (ascites or tumors) (Figure 4B). However, it should be noticed that the response to doxorubicin present some discrepancies between models of the same patient, in both patients (Figure 4B).

Interestingly, 4 PDTO models were established from 4 HGSOC patients with recurrence: patient OV-021 was sampled before the second line of treatment, patients OV-015 and OV-150 before the third line and patient OV-026 before the fourth line. All samples at recurrence were ascitic fluids (Supplementary Table 2). The 4 recurrent PDTO displayed heterogenous response to the 6 drugs tested (Figure 4C), with most of them on the spectrum of intermediate to resistant (Figure 4A), probably due to the impact of the previous treatment lines.

### PDTO HR status determination

Considering the importance of PARPi maintenance treatment for the therapeutic management of ovarian cancers, we focused our attention on the evaluation of the potential use of PDTO for clinical decision making in this regard. It is commonly admitted that the platinum-sensitivity is one of the criteria of eligibility to PARPi. Therefore, the correlation between the PDTO response to carboplatin and to PARPi in our panel of 29 PDTO was evaluated. PDTO response to carboplatin and olaparib displayed a modest correlation (Spearman r = 0.32, p = 0.089) (Figure 5A). Correlation between niraparib and carboplatin, olaparib and niraparib, were also tested and showed statistical significance (Spearman r = 0.57, p = 0.001 and Spearman r = 0.52, p = 0.004, respectively) (Supplementary Figures 5A and B). We then studied the correlation between the PDTO response to olaparib and HRD/HRP status of PDTO determined by the genomic instability score GIScar^8^. However, no correlation was observed between the GIScar analyses and the PDTO response to olaparib (Figure 5B), neither to carboplatin (Supplementary Figure 5C) nor to niraparib (Supplementary Figure 5D).

**Figure 5.**
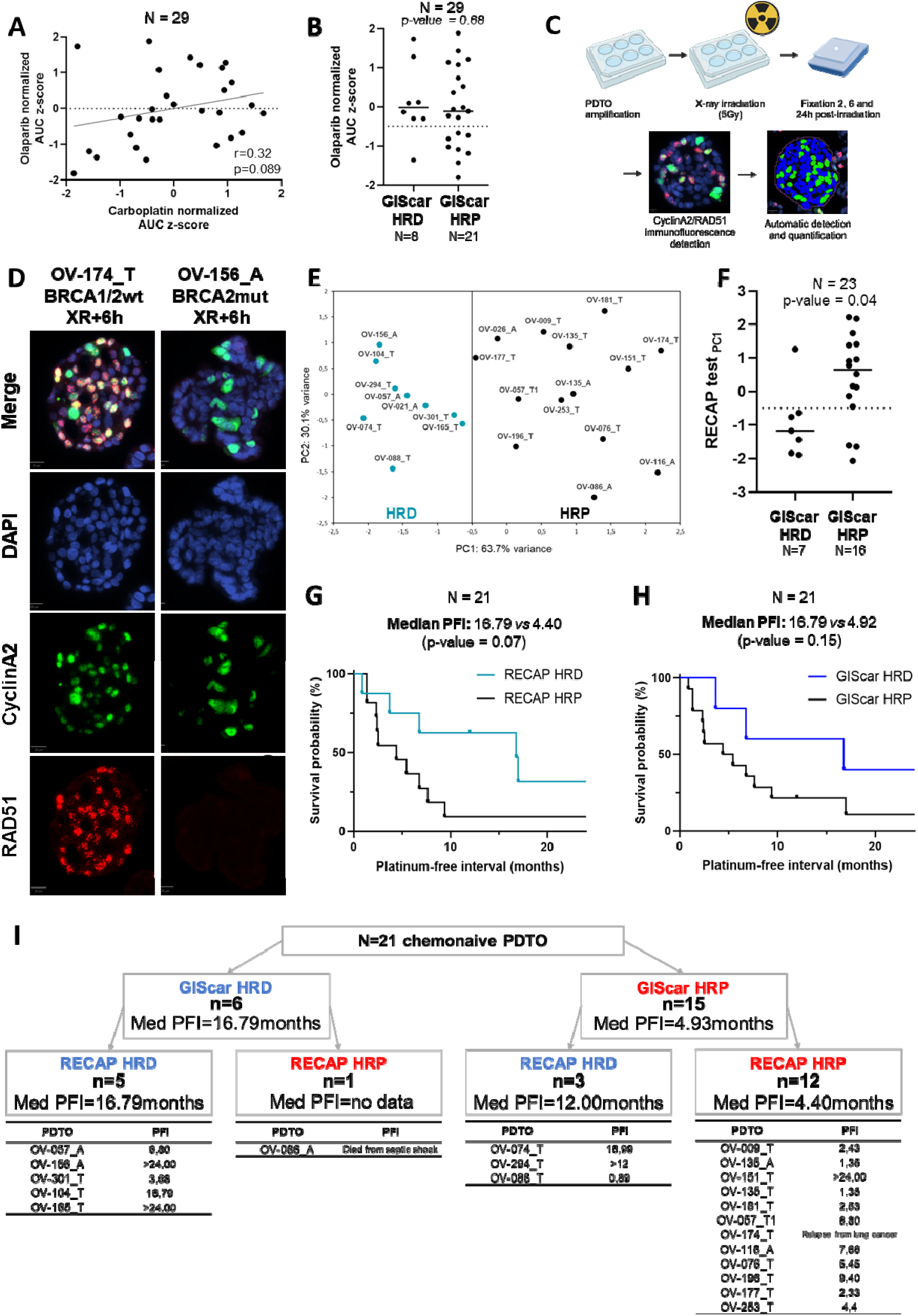
PDTOs can be used to assess homologous recombination deficiency. A. Correlation between the carboplatin and olaparib PDTO response, expressed as normalized AUC z-score (n=29). B. Dot plots comparing PDTO response to olaparib, expressed as normalized AUG z-score, and the genomic Instability score HR status (n=29). Data were analyzed using unpaired two-sided Mann-Whitney test. C. Schematic representation of the RECAP test protocol. D. DAPI staining to visualize nuclei (blue) and immunostaining of cyclinA2 (green) and RAD51 (red) In BRCAI/2-wl (OV-174_T) and BRCA2-mut (OV-156_A) PDTO models. E. Principal component analysis (PCA) of RECAP test results (n=23). A = Ascites-derived PDTO and T = Tumor-derived PDTO. F. Dot plots comparing PDTO RECAP test PC1 and the genomic Instability score HR status (n=23). Data were analyzed using unpaired two-sided Mann-Whitney test G. Kaplan-Meier plot comparing PFI of the RECAP HRD group (n=8) and the RECAP HRP group (n=13). H. Kaplan-Meier plot comparing PFI of the GIScar HRD group (n=6) and the GIScar HRP group (p=15). I. Median PFI of ovarian cancer patients from PDTO were derived depending on the HR status determined by GIScar analysis and RECAP test.

In order to determine another functional assay to identify patients who could benefit from PARPi, we investigated the predictive potential of the RECAP test^27^, and its automated analysis previously set up by our team (Thorel et al., Lab. Invest., in press). Briefly, PDTO were exposed to X-ray and then grown for various times before being embedded in paraffin for immunofluorescence detection of RAD51 in CyclinA2 positive cells (Figure 5C and D). Principal component analyses (PCA) distinguished two populations of PDTO models defined as HRD and HRP using a threshold of −0.5 on the PC1 axis (Figure 5E). This analysis did not allow the establishment of a correlation between HR status and PDTO response to carboplatin, olaparib and niraparib (Supplementary Figure 5E, F and G). However, this RECAP-based HR status was strongly correlated to the standard definition of HR status (genomic instability score, here defined by GISCar score) (p-value = 0.04) (Figure 5F). Otherwise, the analysis of the PFI in this set of patients (after exclusion of 2 PDTO derived from samples obtained at recurrence) showed that RECAP HRD group (n=8) had a median PFI of 16.79 months vs 4.40 months in the RECAP HRP group (n=13) (p-value = 0.07) (Figure 5G). Similar values were obtained with GIScar score with a median PFI of 16.79 months vs 4.92 months (p-value = 0.15) (Figure 5H). These analyses have also been performed on HGSOC subgroup (n=17) but are not conclusive (Supplementary Figure 5H and I). In addition, the RECAP test could bring supplementary information in the context in which genomic analyses lead to the conclusion of an HRP status (Figure 5H). Indeed, in this arm (GIScar HRP, n=15), our analysis of RECAP test reclassified 3 GIScar HRP as HRD. Interestingly, 2 of these 3 patients had a PFI superior to 12 months (median PFI = 12 months in GIScar HRP/RECAP HRD subgroup vs 4.40 months in GIScar HRP/RECAP HRP subgroup).

## Discussion

We successfully established 37 long-term PDTO models from 31 patients with ovarian cancer. The majority of the PDTO of the panel was derived from patient diagnosed with HGSOC, in accordance with the high frequency of this subtype observed among ovarian cancer patients (70% of all epithelial ovarian cancers)^2^. We were also able to produce PDTO models from rarer subtypes including OCCC, CS, MC and EM (for a total of 11 PDTO representing 8 patients). To our knowledge, this is the largest panel of PTDO from OCCC and CS although other teams had also been able to generate PDTO from such subtypes^21,22,28–30^. Knowing that most of these subtypes (especially OCCC and CS) are resistant to platinum-based chemotherapies, such preclinical models could be relevant to develop new therapeutic strategies for these tumors with few treatment options.

In this study, we achieve a long-term ovarian PDTO establishment rate (number of passages above 8) of 13.8%, showing that only a small proportion of ovarian cancers patient could currently benefit from the PDTO-based precision medicine. However, this rate of establishment was difficult to compare with other studies since there is no consensus on the definition of a successful establishment of PDTO. In fact, many studies used different criteria of establishment and different definitions of long-term culture^25^. Overall, the success rate of long-term establishment (often defined as a number of passages above 4) varies from 18% to 53%, depending on the study considered^17,24,28,29^. Senkowski et al. were able to improve their rate of establishment by systematically testing two different culture media to achieve long-term culture of PDTO from HGSOC in 53% of the cases, highlighting the importance of culture conditions^24^. In order to represent a larger proportion of ovarian cancer patients, it would be necessary to work on earlier passages of PDTO. Indeed, as demonstrated by Bi et al., the use of PDTO during the first passages allowed to perform drug sensitivity assay for 83% of the tumor samples^30^.

We characterized our PDTO panel with a set of tumor markers routinely used in ovarian cancers diagnosis and PDTO showed a strong ability to reflect the features of their sample of origin in accordance with their histotypes. Genetic analyses also demonstrated a tight correlation between the CNV of the original tumor and that of paired PDTO. Thus, these analyses showed that PDTO not only recapitulate inter-patient heterogeneity, but also that our PDTO models are matched with their tumor of origin. Interestingly, intra-patient heterogeneity was retained as well, since PDTO from both tumor and ascitic samples of 2 patients (OV-057 and OV-135) maintained their original sample CNV profile. At the transcriptomics level, despite a strong correlation between the gene expression profile of the tumor sample and the paired PDTO, clustering showed that tumors were still closer to each other than to their PDTO. An alternative approach using non-negative matrix factorization allowed us to only consider the cancer component of paired samples and revealed that the PDTO recapitulate the transcriptomic profile of their tumor of origin in our relative long-term culture.

To assess whether ovarian PDTOs could predict patient response to first-line chemotherapy, only chemonaive PDTO were selected, since PDTO from previously treated patients might have gained alterations that could introduce bias in response to treatments. We were able to match the PDTO response to carboplatin and the PFI of the patients following the first line treatment. Out of 9 tested patients, 2 displayed a difference between the clinical response and the paired PDTO sensitivity to carboplatin, whether PDTO were derived from a carcinomatosis node or from ascitic sample. The medical history of these patients showed that, although they did not undergo complete surgical resection at the initial clinical management, both patients responded well to the first cycles of chemotherapy, as evidenced by the drastic decrease in CA-125 and the partial response assessed by mid-line RECIST. These observations suggested a response from the whole tumor mass, in accordance with the observations made on their PDTO. This shows that PDTO can predict tumor cell response to first-line carboplatin. However, in some cases of highly aggressive tumors or tumors showing a great plasticity, rapid recurrence may be observed despite a good initial clinical response, itself correlated with the PDTO response. For the concerned patients, the recurrence was detected in a mediastinal lesion (OV-135) or in a periaortic lymph node (OV-301), which may also suggest a sampling bias representing one of the limitations of these PDTO-based functional assays. Nevertheless, this limitation is also observed in all types of predictive assays (genomic, transcriptomic or functional assays), and the tumor heterogeneity in some patients may constitute an obstacle to the use of these functional assays for predictive purposes.

The main limitations of this study are the low establishment rate and the delay between the tumor/ascites sampling and the results of the predictive functional assay, making it currently not suitable in a context of clinical routine. In this study, the main objective was to assess the feasibility of establishing ovarian PDTO and to study their representativity of the patient at a genomic, transcriptomic and functional level in order to evaluate their predictive potential. We therefore established long-term PDTO, as they have the advantage of being stable and widely amplifiable for basic and translational applications. Regarding PDTO use for precision medicine, it becomes necessary to work on early passages despite small amount of PDTO, to be able to perform the functional assay rapidly and for most of the patients (around 90% according to Bi and al.^30^).

In this study, numerous PDTO models were established from ascitic samples (11/37) an integrative biological material of particular interest since ascites is present in 1/3 of ovarian cancer at the initial diagnosis and in a large majority of cases at the recurrence^31^. It would therefore enable a potential longitudinal assessment of patient response to treatment. Moreover, PDTO from ascites would also be usable for predicting response to treatment at recurrence, when surgery is rarely considered, thereby making difficult the obtention of tumor cells. Herein, we collected paired tumor/ascitic samples from two HGSOC patients. Interestingly, paired PDTO from these samples displayed a similar sensitivity to all the drugs tested with the exception of doxorubicin. More paired PDTO models derived at the same or different time of the clinical course are needed to conclude about the predictive potential of ascites-derived PDTO, as initiated by Arias-Diaz et al^32^. Future studies will also need to investigate the ability of PDTO established from post-treatment samples (second and later lines of therapy) to provide effective guidance for therapeutic decisions in the management of patients in more complicated clinical situations, with fewer therapeutic options available, especially in the context of platinum-resistance recurrence.

While PDTO response to carboplatin showed a tight correlation with paclitaxel response, modest correlation were found between carboplatin and the two PARPi tested (olaparib and niraparib), in line with platinum sensitivity being suggested as a biomarker for predicting response to PARPi^33^. However, the PDTO response to PARPi or carboplatin was not correlated to HRD status determined either by GIS or by our novel approach using the RECAP test. On the other hand, this RECAP test classification of HRP and HRD PDTO was closely correlated to the GIScar scoring as well as to the patient clinical response to first line chemotherapy defined by the PFI. Unfortunately, only a very small number of patients enrolled in this study were prescribed PARPi during their clinical management, making it impossible to establish a correlation between functional assays and the patient clinical response to PARPi. Therefore, since the GIScar scoring is closely correlated with the patient response to olaparib^8^, it is the best indicator of HRD classification available so far. These results suggest that direct exposure to olaparib is less effective than the RECAP test in predicting patient response to carboplatin. However, the RECAP test may also have its limitations (RAD51 foci cannot reflect HRD cells defective in steps down-stream of RAD51 foci formation or drug efflux), and it remains worthwhile pursuing studies to optimize the modalities of the functional assay based on the direct exposure of PDTO to PARPi to correlate with the clinical response.

Furthermore, our study suggests that testing the functionality of the HR pathway based on the RECAP test could offer superior performance to the genomic test. Indeed, within the group of patients identified as HRP by GIScar, the patients confirmed HRP by the RECAP test showed a median PFI of 4.4 months while the patients identified as HRD by the RECAP test had a median PFI of 12 months (Figure 5I). In this this latter group, 1 of the 3 patients concerned appears to be resistant to treatment (PFI = 0.89). However, the other 2 patients have PFIs of 16.98 and >12 months, strongly suggesting that the RECAP test has classified them correctly. Thus, RECAP test could identify HRD patients who had not been previously identified using genomics and enabling them to benefit from PARPi.

In conclusion, our study shows the feasibility and the interest of generating ovarian PDTO of different histotypes to predict response of first line treatment, and highlights the potential of functional assays to complement the genomic tests currently used for PARPi prescription. Although our cohort has its limitations (small size and heterogeneity of clinical parameters), it is one of the largest collections described to date. This study also highlights the many challenges that still need to be met before the routine use of these assays in clinical oncology. This includes the necessity of shortening the time needed to assess response to treatment, reducing the quantity of material required to perform these functional assays, and increasing the success rate of PDTO culture. It is thus essential that future investigations focus on the application of functional assays as soon as possible, ideally just after the formation of the first PDTO (early passages). This will involve to determine their ability to summarize the characteristics and heterogeneity of the tumors from which they are derived, and will require the development of miniaturization and standardization of functional assays. It will also be necessary to compare PDTO results to functional assays whether they are used in early passage or after long-term establishment. The presence of stromal cells in the first passages is also likely to modify response to treatment. This may be an advantage, but must be monitored to ensure sufficient robustness to enable these tests to be applied to a precision medicine approach in ovarian cancers. All the work being carried out by the community on these burning issues is opening up both interesting prospects and new questions, but it is only if they are resolved that PDTOs will find their full potential in the management of these cancers.

## Methods

### Patient-Derived Tumor organoid establishment

#### Ethical considerations

Fresh tumoral tissue and ascitic fluids from ovarian cancers were collected from patients treated at the Comprehensive Cancer Center Francois Baclesse (Unicancer Center, Normandy). Informed consent forms were signed by all patients and were obtained either by the Biological Resources Center ‘OvaRessources’, which has received NF 96 900 accreditation (N° 2016/72860.1) or in the context of the ‘OVAREX’ clinical trial (N°ID-RCB: 2018-A02152-53, NCT03831230)^34^, in accordance with ethical committee and European law. Clinical, treatment and histopathological details were extracted from patient charts. A medical pathologist analyzed all samples.

#### Processing of samples

Samples were processed as previously described^14^. Briefly, tumor tissue was cut into 4 mm^3^ pieces. One piece was fixed in 3% paraformaldehyde for paraffin embedding and subsequent histopathological/immunochemistry analyses, two pieces were snap frozen in FlashFreeze (Milestone) and stored at −150°C for DNA/RNA extractions, and another piece was processed to establish PDTO. Tumor sample dissociation was performed using the Tumor Dissociation human kit and a gentleMACS Dissociator according to the manufacturer’s instructions (Miltenyi Biotec). Sterile tumor ascitic samples were centrifuged (430 g for 5 min). Pellets containing cells were resuspended in 20 mL of RPMI 1640 medium (Fisher Scientific) supplemented with 10 UI/mL penicillin, 10 µg/mL streptomycin (Fisher Scientific) and 1% Bovine Serum Albumin (BSA) (Sigma). Suspensions were strained successively in 300 µm and 50 µm filters (Endecotts). Remaining cells or aggregates were digested in 2 mL of TrypLE Express (Fisher Scientific) at 37°C up to min. Dissociated cells from tumors and ascites were collected in organoid basal medium [OBM: Advanced DMEM (Fisher Scientific), 10 UI/mL penicillin, 10 µg/mL streptomycin, 1% GlutaMAX-1 (Fisher Scientific)] and pelleted (430 g for 5 min). 10 000-20 000 cells were resuspended in organoid culture medium (OBM supplemented with B27 (Fischer Scientific, 200 µL/mL), N-Acetyl-L-cysteine (Sigma, 1.25 mM), EGF (Miltenyi, 50 ng/mL), FGF-10 (Peprotech, 20 ng/mL), FgF-basic (Miltenyi, 1 ng/mL), A-83-01 (Peprotech, 500 nM), Y27632 (Selleckchem, 10 µM), SB202190 (Peprotech, 1 µM), Nicotinamide (Sigma, 10 mM), PGE2 (Sigma, 1 µM), Primocin (InvivoGen, 100 µg/mL), cultrex HA-R-Spondin-1-Fc 293T (AMS Bio, 50% V/V) and Cultrex L-WRN (AMS Bio, 10% V/V), mixed with 70% Cultrex Reduced Growth Factor Basement Membrane Extract, Type 2 (BME2) and seeded in a pre-warmed 24-well plate (Eppendorf). After polymerization (37°C, 5% CO2, 15 min), each drop was immersed with 500 µL of organoid culture medium. Medium was exchanged twice a week. Once harvested with OBM supplemented with 1% BSA (OBM-BSA), PDTO were dissociated using TrypLE Express (37°C up to 10 min). Isolated cells were seeded or biobanked (Coolcell, −80°C) in 500 µL of Recovery cell culture freezing medium (Fisher Scientific) for future use.

#### PDTO culture

When PDTO reached around 150 µm in diameter, they were collected with cold OBM-BSA, centrifuged at 200 g for 2 min and incubated with TrypLE Express for up to 15 min at 37°C. After dissociation, cells were centrifuged at 430 g for 5 min, suspended in organoid culture medium, counted and plated at 10 000 cells per 50 µl drop of 70% BME2 in pre-warmed 24-well plates. Plates were transferred to a humidified 37°C/5% CO2 incubator. Cryovials were prepared at regular intervals by dissociating and resuspending PDTO in Recovery Cell Culture Freezing Medium (Gibco), then placed in a cell freezing container (Coolcell) at −80°C and biobanked at −150°C on the next day. PDTO lines were authenticated by comparison of their short tandem repeat (STR) profiles with that of tumor sample of origin (Microsynth).

### Characterization

#### Histology and immunohistochemistry

Tissue and PDTO were fixed in 3% paraformaldehyde overnight. After the embedding of PDTO in 2% agarose, samples were dehydrated, paraffin embedded, and sectioned before standard hematoxylin and eosin staining (H&E). Automated immunohistochemistry using a Ventana Discovery Ultra (Roche) was performed on 4 µm-thick paraffin sections. Slides were deparaffinized with EZPrep buffer and epitopes were unmasked by 56 min of high-temperature treatment in CC1 EDTA buffer. Sections were incubated for 40 min at 37°C with an anti PAX8 (ab191870, Abcam, 1/500), p53 (ab16665, Abcam, 1/100), HNF1β (ab213149, Abcam, 1/2000) or Ki67 antibody (NCL-Ki67p, Novocastra, 1/500). Secondary antibody (Omnimap Rabbit HRP; Ventana Medical System Inc., Tucson, AZ, USA) was incubated for 16 min at room temperature. Immunodetection performed without the primary antibody was used as control. After washes, the staining was performed with DAB (3, 3’-diaminobenzidine) and sections were counterstained with hematoxylin using Ventana reagents according to the manufacturer’s protocol. Stained slides were then digitized with a 20X magnification using the Vectra Polaris slide scanner (Akoya Biosciences).

### Genomic and transcriptomic analyses

#### DNA and RNA extraction

DNA and RNA extractions were performed using the NucleoSpin RNA and tissue kits according to the manufacturer protocol (Macherey–Nagel). After extraction DNA and RNA samples were stored at −80°C.

#### Low pass WGS

Low Pass WGS was performed on DNA samples. Library was prepared using Illumina DNA PCR-Free Prep (Illumina) starting from 500 ng of DNA input (except for 6 samples: 93, 356, 384, 401, 430 and 483ng). Libraries were pooled all together to be sequenced on one flowcell. 150 bp paired-end sequencing of the samples was performed on the NovaSeq 6000 (Illumina). Raw reads were mapped to the human reference genome (GRCh38) using the Burrows-Wheeler Aligner (BWA) with MEM algorithm (version 0.7.17). Read duplicates were removed by Picard MarkDuplicates (version 2.27.5). Read count was performed by HMMcopy (version 0.1.1) with a bin (non-overlapping window) of 50 kb. Copy number alteration (CNA) identification and tumor fraction estimation were performed using ichorCNA (version 0.3.2). For data visualization, R/Bioconductor packages karyoploteR (version 1.16.0) and copynumber (version 1.30.0) were used.

#### Transcriptome

Starting from 4 µl of total RNA we add 1 µl of ERCC spike-in control. Library generation is then initiated by oligo dT priming, from total RNA (200 ng RNA input in 4 µl) following Lexogen QUANTSEQ FWD + UMI protocol. After first strand synthesis the RNA is degraded and second strand synthesis is initiated by random priming and a DNA polymerase. At this step Unique Molecular Identifiers (UMIs) are introduced allowing the elimination of PCR duplicates during the analysis. After obtaining the double stranded cDNA library, the library is purified with magnetic beads and amplified. During the library amplification, the barcodes and sequences required for cluster generation (index i7 in 3’ and index i5 in 5’) are introduced due to Illumina-compatible linker sequences with 14 cycles of amplification. The final library is purified and checked on a High sensitivity DNA chip to be controlled on the Agilent bioanalyzer 2100. Each library is pooled equimolarly and the final pool is also controlled on Agilent bioanalyzer 2100 and sequenced on NovaSeq 6000 (Illumina) (100 bp single-end).

#### Data analysis for transcriptomic data

To eliminate poor quality regions and poly(A) of the reads, we used the fastp program. We used quality score threshold of 20 and removed the reads shorter than 25 pb. The read alignments were performed using the STAR program with the human genome reference (GRCh38) and the reference gene annotations (Ensembl). The UMI (Unique Molecular Index) allowed to reduce errors and quantitative PCR bias using fastp and umi-tools. Based on read alignments, we counted the number of molecules by gene using FeatureCount. Others programs were performed for the quality control of reads and for the workflow as qualimap, fastp, FastQC and MultiQC. Differential Gene Expression of RNA-seq was performed with R/Bioconductor package DESeq2.

#### NMF on transcriptomic data

Non-negative Matrix Factorization (NMF) was applied using the butchR package. We started from a matrix of normalized expressed genes (13 401 expressed genes) and applied variance stabilizing transformation (VST). We started to apply NMF on a subset of the VST matrix subsetting to Organoïd samples only. For each analysis different rank (k) were tested (from 2 to 18). Then we projected the NMF factors on all samples (tumors and organoïds) using the LCD function from the R YAPSA package using as input the count matrix (VST) and the W matrix from the NMF applied on organoïd samples. Given the NMF on all samples, we plotted UMAP projection using ggplot and computed distance between paired samples (organoid/tumor) based on the UMAP. To choose the best rank, we choose the one that minize the sum of distances between paired samples (organoid/tumor). We then fixed the rank to k=12.

### Evaluation of response to treatments

#### Drugs

After reconstitution in saline solution, doxorubicin (Teva) and gemcitabine (Sandoz) were stored at 4°C, paclitaxel and carboplatin (Accord) were stored at room temperature. Olaparib and niraparib (Medchemexpress) were diluted in DMSO and stored as stock solution at −80°C.

#### PDTO treatment

When PDTO reached the size of 75-150 µm in diameter, PDTO were collected with cold OBM-BSA and centrifugated at 200 g for 2 min. PDTO were resuspended in organoid treatment medium (organoid culture medium lacking primocin, Y-27632 and N-acetylcysteine) and counted. For treatment in 96-well plates, PDTO were resuspended in 2% BME2/organoid treatment medium and 200 PDTO per well were seeded in 100 μL volume in a previously coated (1:1 organoid treatment medium/BME2) white clear bottom 96-well plates (Greiner). For treatment in 384-well plates, PDTO were resuspended in 10% BME2/organoid treatment medium and 50 PDTO per well were seeded in 20 μL in black ULA clear bottom 384-well plates. Thirty minutes later, PDTO were exposed to treatments and PDTO morphology was monitored using CellDiscoverer 7 (Zeiss). One week later, ATP levels were measured by CellTiter-Glo 3D assay (Promega) and luminescence was quantified using GloMax (Promega). Cell viability values were normalized to control and treatment sensitivity was expressed as the average of at least two independent biological replicates. Positive control (Staurosporine 10µM) was used to calculate a z-score as followed:

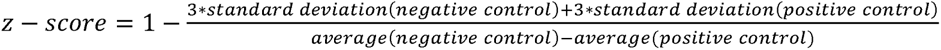

An experiment with a z-score under 0.4 was considered as non-exploitable. Viability curves were designed using GraphPad Prism software (version 9.2.0). The area under the curve (AUC) was computed for each PDTO model. Normalized AUC z-scores were calculated by subtracting the mean and divided the results by the standard-deviation of the AUC compared to each AUC value.

##### Genomic instability scoring of PDTO

In order to assess PDTOs homologous recombination (RH) status, PDTOs were sequenced with a 127-genes panel including 15 HR genes (*BRCA1, BRCA2, ATM, BARD1, BRIP1, CDK12, CHEK1, CHEK2, FANCL, PALB2, PPP2R2A, RAD51B, RAD51C, RAD51D, RAD54L*). The sequencing data were then be used to determine a genomic instability score (GIS) as described by Leman et al^8^. Briefly, this genomic instability score was based on a first analysis by a CNVkit pipeline v0.9.7^35^. From the CNVkit data outcome, three scores of instability were calculated: the number of large genomic events, the structural instability score, and the allelic imbalance. Then, these scores were used in the logistic model to compute a mathematical esperance that the tumor is HR-proficient (HRP) or HR-deficient (HRD), with a value tending to 0 for HRP tumors and a value tending to 1 for HRD tumors. The decision-making threshold between HRP and HRD tumors was 0.48. However, this threshold was defined on FFPE tumor tissue and at the time of this work we could not propose an optimal threshold for PTDO. We defined so a third category HRDmid corresponding to PTDO with a score at more and less 0.25 than the threshold of FFPE tumor tissues (0.48), i.e. a range score of [0.23 – 0.73].

### RECAP test

#### PDTO irradiation

To evaluate the homologous recombination deficiency status, we used the RECAP test described by Naipal and al.^27^. PDTO were grown in BME-2 domes in 6-well plates. When PDTO reached the size of 75-150 µm in diameter the medium was collected and conserved at 37°C. Two ml of OBM were added to each well (including control plate) before PDTO were irradiated at 5Gy (X-Ray source, 130kV, 5mA) in a CellRad X-irradiation system. After irradiation, the medium was discarded and replaced by the previously conserved warm organoid culture medium. Following incubation of the plates for 2h, 6h or 24h, PDTO were collected, washed with OBM-BSA and fixed in 3% PFA for at least 3 hours at +4°C. PFA was then discarded and the pellet was embedded in agarose, dehydrated and paraffin-embedded (Autostainer XL, Leica).

#### PDTO staining

Immunofluorescence staining of RAD51 (ab133534, Abcam) and Cyclin A2 (ab211736, Abcam) were done on 4 µm sections of PDTO (microtome Leica) using Ventana Discovery Ultra (Roche). Microscope slides with PDTO sections were deparaffinized with EZPrep buffer and the antigenic sites were unmasked with EDTA. The endogen peroxidase inhibitor (Roche) was used to limit to background noise. 100 µl of primary antibody previously diluted (Primary Antibody Diluant, Clinisciences) were added to each slide and incubated at 37°C for 40 min (1/100 000 for RAD51 and 1/1 000 for Cyclin A2). Then, after washing, the second antibody (Omnimap Rabbit) was incubated at 37°C for 16 min. Finally, the DISCOVERY Rhodamin (Roche) and the DISCOVERY FITC (Roche) were added for RAD51 and Cyclin A2, respectively. The slides were subsequently stained for DAPI (Roche). At the end of the procedure, the slides were washed with detergent to eliminate the LCS oil layer and were hydrophilic-mounted with Fluoromount-G (CliniSciences). The slides were protected from light until digital acquisition.

#### Digital acquisition

Specimens of PDTO sections were digitized with an Olympus VS120 scanner equipped with a 40x objective (N.A 0.95), a LED illumination (Lumencor Spectra X 7 LED), a single multiband filter DAPI (emission: 455nm; exposure time: 5ms), FITC (518nm; 20ms) and CY3 (565nm; 10ms) filters from Olympus and a CMOS camera from Hamamatsu (Orca Flash 4.0). The whole slide images (WSI) were recorded with Extended Focal Imaging (EFI) technology applied over a height of 4 µm (Z-range) and an acquisition of 5 slices per field (0.84 µm Z-spacing).

#### Quantification processing

On each slide, the PDTO were isolated automatically by detection with Qupath program and the extension Cellpose^36,37^. For each individual PDTO, nuclei detection was assessed on the DAPI channel and the surface occupied by nuclei was processed using extension Stardist of Qupath program^38^. For the positive nuclei detection of Cyclin A2, two Gaussian filters of different sizes were applied on the FITC channel. Only the positive pixels obtained after the difference between the two filtered images were kept to obtain a binary image with the positive Cyclin A2 areas. Then a geodesic reconstruction was applied between the images of nuclei and Cyclin A2 to identify only positive nuclei with a Cyclin A2 active. Finally, for the detection of RAD51 foci, the CY3 channel was used. RAD51 foci were detected thanks to a Laplacian filter after a Gaussian operation, in the scikit-image python library^39,40^. Only bright foci on dark backgrounds are detected. For each nucleus of each organoid, only the RAD51 foci inside the CyA2-positive nuclei were kept and computed. ***Gaussian Mixture Model approach.*** A Gaussian Mixture Model (GMM)^41^ was used to determine the minimum threshold of RAD51 foci in proliferative cells (Cyclin A2-positive nuclei), making it possible to differentiate the basal level of HR DNA repair from irradiation-induced HR DNA repair. Ultimately one Gaussian function was found but it allowed the smoothing of the histograms. Three different indexes were described using the gaussian approach: the damage index (DI) equal to the mean of the control gaussian, the radio-induced damage index (RDI) the difference between the mean of the 2 hours gaussian and the mean of the control gaussian, and the reparation index (RI) the difference between the mean of the 24 hours gaussian and the mean of the 2 hours gaussian. The indexes were computed for each PDTO model and Principal Component Analysis (PCA) was applied to keep only principal components (PC) with maximum variance^42^. A cutoff value of −0.5 was used on the PC1 axis to categorize the homologous recombination status, with the negative value characterizing HRD status and the positive values HRP ones.

### Statistics

Quantitative variables were compared with unpaired two-tailed Mann-Whitney test using GraphPad Prism. Correlations between gene expression levels of PDTO and tumor were calculated using Pearson’s R. Survival were calculated using the Kaplan-Meier method, and survival distributions were compared using the Log-Rank test. P values <0.05 were considered significant. Definitions of n and details of statistical analyses are provided in relevant figure legends.

## Acknowledgements

This work is supported by Cancéropôle Nord-Ouest, Ligue contre le Cancer (Calvados’s commitee), Fondation de l’Avenir (#AP-RM-19-020), the “Fondation ARC pour la recherche sur le cancer” (#PJA20191209649). “ORGATHEREX” project was co-funded by the Normandy County Council, the European Union within the framework of the Operational Programme ERDF/ESF 2014-2020 which was conducted as part of the planning contract 2015-2020 between the French State and the Normandy Region. The funders had no participation in study design, data management, or publication management.

The ORGAPRED core facility ‘Tumor organoids for research and predicting response to treatment’ from the PLATON services unit (University of Caen Normandie) and research associated programs “ORGAPRED”, “PLATONUS ONE”, “POLARIS”, “Equip’Innov Caen 2022 – PLATONUS-ONE” and “Normandie Recherche Plateformes 2023 – PLATONUS 2023” projects were supported by the Normandy County Council and the European Union within the framework of the Operational Programme ERDF/ESF 2014-2020) and by the French state as part of the planning contract 2015-2020 between the State and the Lower Normandie Region. We thank the donors, “Vaincrabe” and “Contre le Cancer, J’y Vais” associations, the Lions clubs of Normandy, and the other foundations for their support of the projects carried out by our teams on PDTO. We are grateful to Inserm, University of Caen Normandy and the Comprehensive Cancer Center François Baclesse for their support in the implementation of these activities. LT has been successively recipient from a doctoral scholarship from French Ministry of Research and Higher Education and from Comprehensive Cancer Center F. Baclesse. PMM was supported by the Cancer Institut Thématique Multi-Organisme of the French National Alliance for Life and Health Sciences (AVIESAN) Plan Cancer 2014-2019 (doctoral grant).

The authors thank Nicolas Goardon, Angelina Legros and Fabrice Guichard from SéSAME facility (Comprehensive Cancer Centre F. Baclesse, Caen, France) who provided technical support for NGS analyses. We also thank Benoît Goudergues (CRB OvaRessource, Services Unit PLATON, Université de Caen Normandie and Comprehensive Cancer Center François Baclesse) for his helpful technical support.

## Data availability

CNA and RNA-seq data will be made available to the community (on GEO or equivalent) upon publication. During the review process, these data will be made available to the reviewers and editor upon request. All other data supporting the findings of this study are available from the corresponding authors on reasonable request. All materials will be available upon request through a material transfer agreement.

## Supplementary information

**Supplementary Figure 1.**
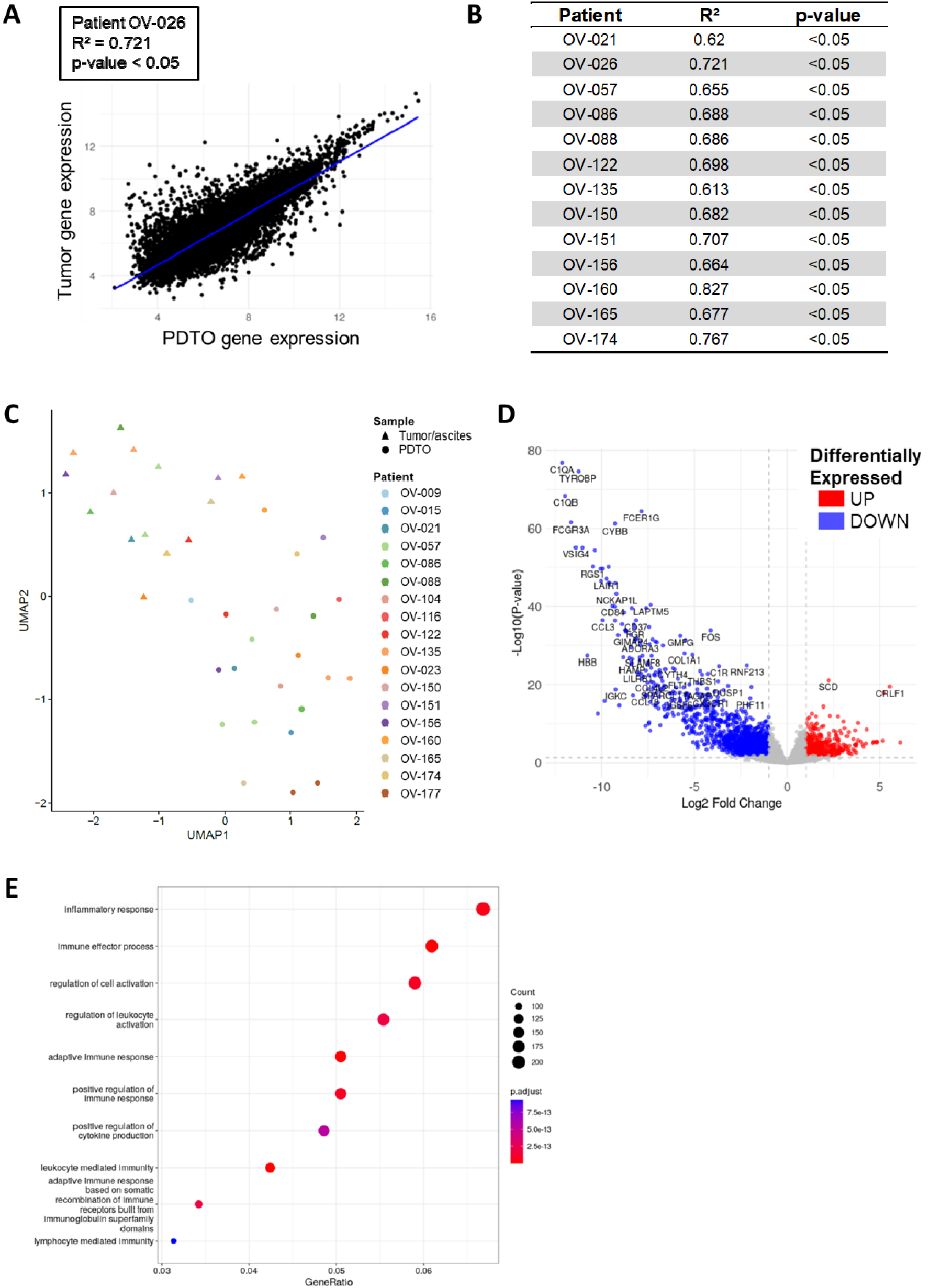
Transcriptomics analysis distinguishes tumor from PDTO samples. A. Scatter plot of the tumor and paired PDTO gene expression for the patient OV-023 with the Spearman’s correlation. B. Spearman’s correlation coefficients of the tumor and paired PDTO gene expression for each patient. C. Distribution of tumor samples and paired PDTO on Uniform Manifold Approximation and Projection (UMAP) clusters based on gene expression profile. D. Volcano plot of differentially expressed genes between tumor samples and paired PDTO. Up- and down-regulated genes are represented in red and blue, respectively. E. Functional enrichment analysis of biological processes (BP) by Over-Representation Analysis (ORA) of differentially expressed genes between tumor samples and paired PDTO.

**Supplementary Figure 2.**
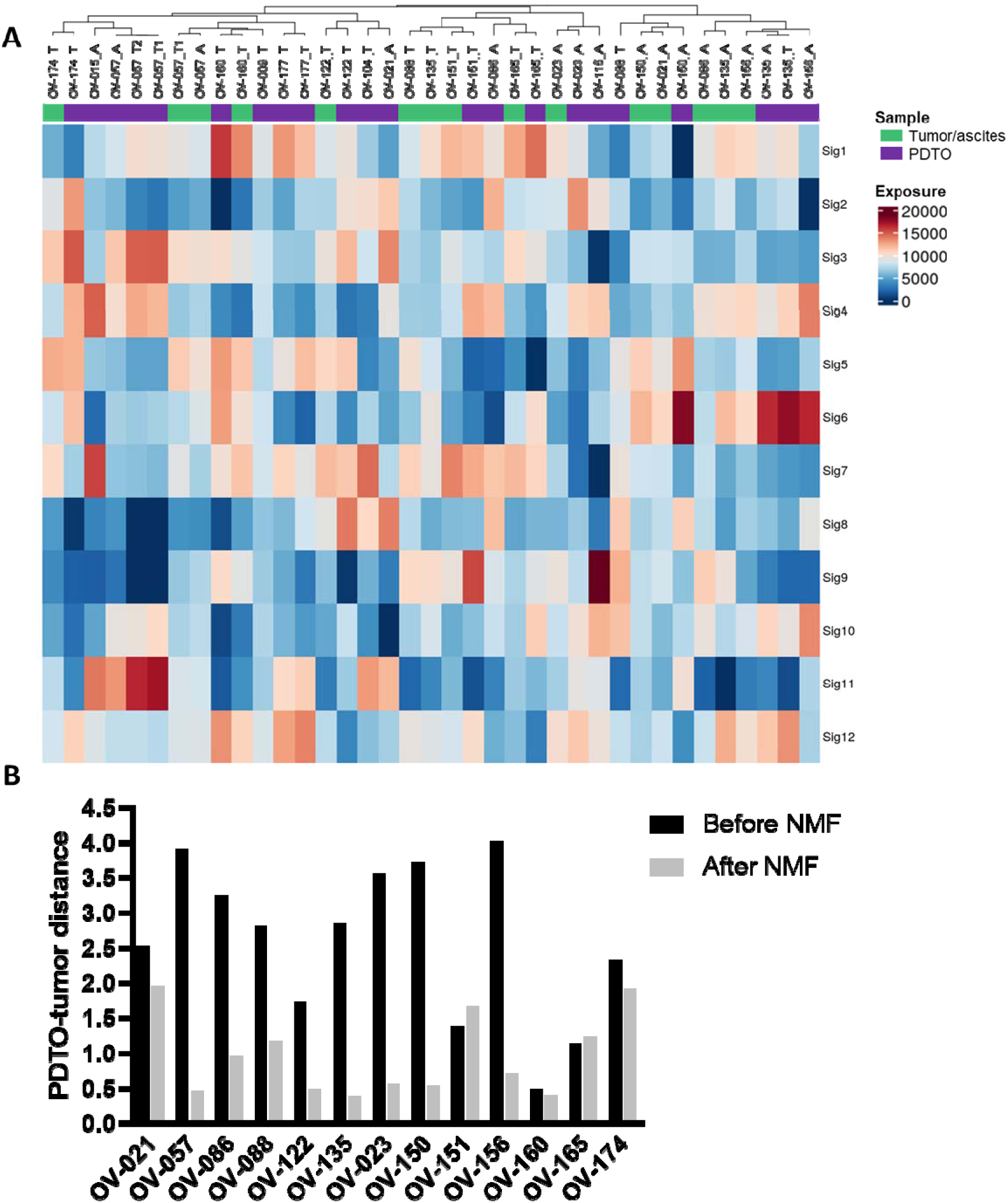
Data dimension reduction brings gene expression profiles of tumors closer to those of their paired PDTO. A. Hierarchical clustering and heatmap of exposure values of PDTO and tumor samples, based on non-negative matrix factorization (k=12) of RNA-seq expression data. A = Ascites-derived PDTO and T = Tumor–derived PDTO. B. Histogram showing distribution of euclidean distances between tumor samples and their paired PDTO using their UMAP coordinates, before and after performing NMF(k=12).

**Supplementary Figure 3.**
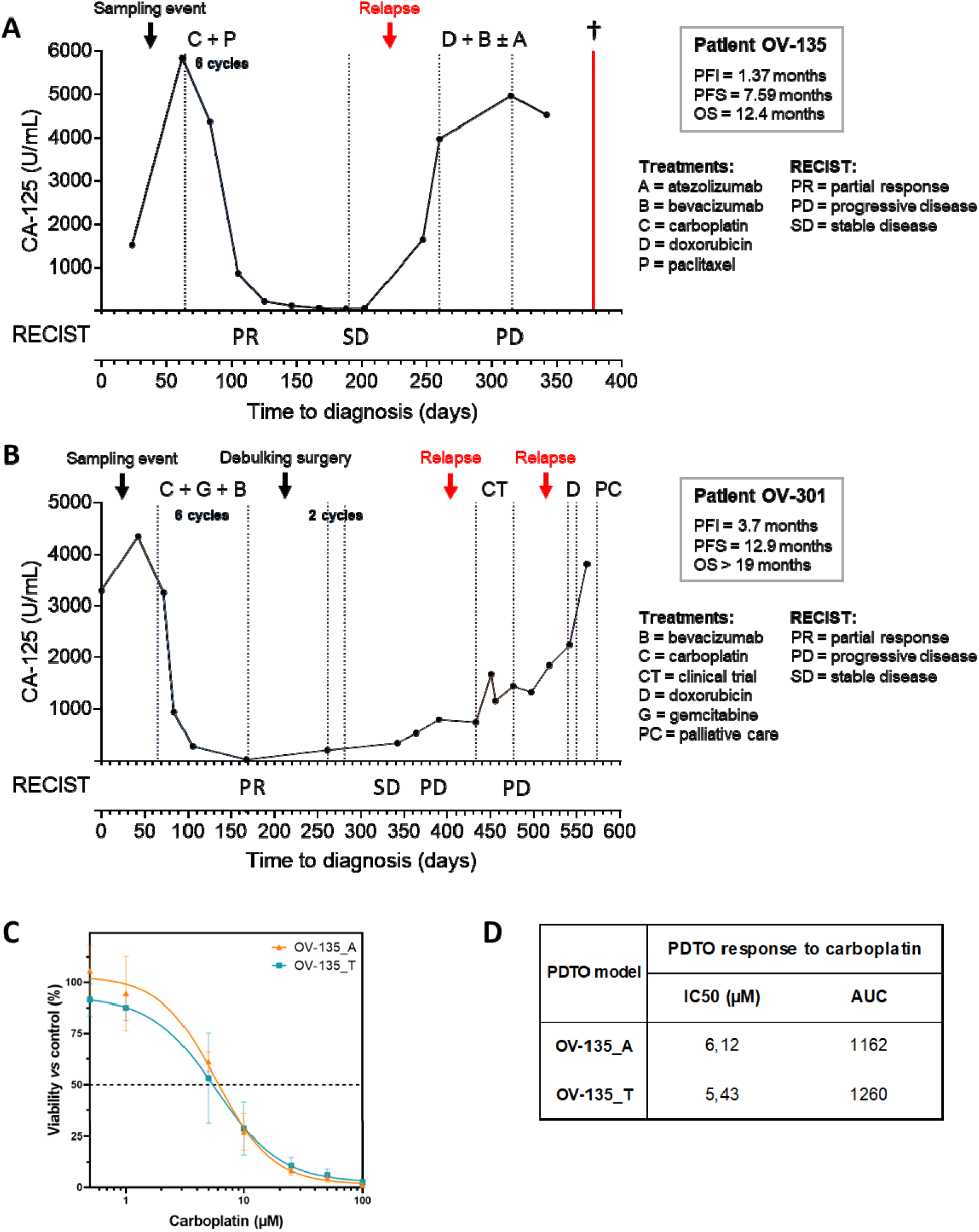
Detailed clinical response of patient OV-135 and OV-301. A. Patient OV-135 response to 1^st^ and 2^nd^ line treatment over time, expressed as serum level of Cancer Antigen 125 (CA-125), RECIST evaluation as well as PFI, PFS and OS. B. Patient OV-301 response to 1^st^, 2^nd^ line and 3^rd^ line treatment over time, expressed as serum level of Cancer Antigen 125 (CA-125), RECIST evaluation as well as PFI, PFS and OS. C. Dose-response curves of the 2 PDTO models OV-135_A and OV-135_T to carboplatin. Each curve is the representative of two independent experiments. D. Summary of the IC50 (µM) and AUC obtained in B.

**Supplementary Figure 4.**
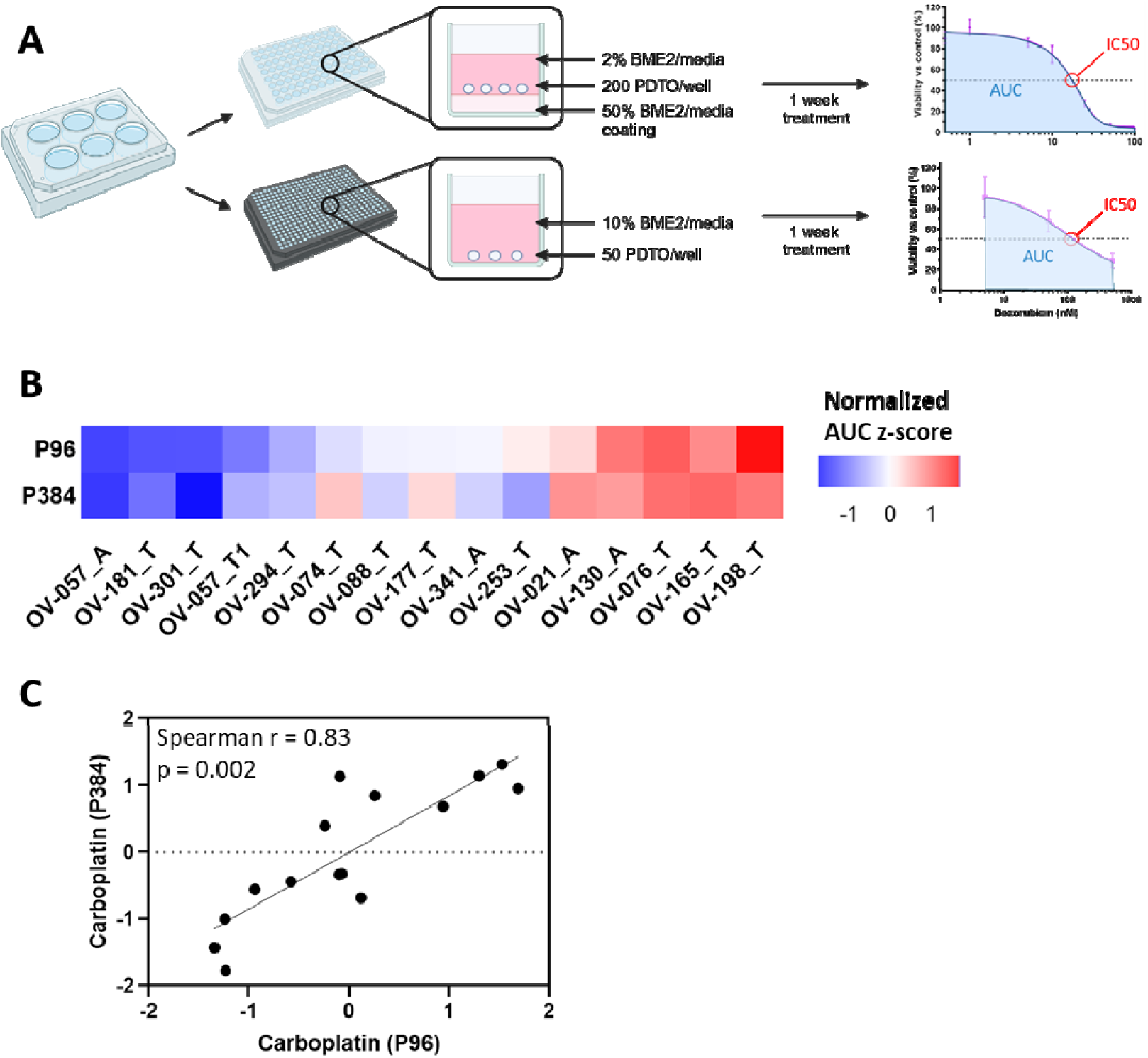
PDTO treatment protocol adaptation from 96-well plate to 384-well plate. A. Schematic representation of PDTO treatment protocols in 96- or 384-well plate. B. Heatmap comparing the response of PDTO to carboplatin, expressed as normalized AUC z-score, in 96- and in 384-well plates (n=15). A = Ascites-derived PDTO and T = Tumor–derived PDTO. C. Scatter plot of the response of PDTO to carboplatin, expressed as normalized AUC z-score, in 96- and in 384-well plate (n=15) with the Spearman’s correlation.

**Supplementary Figure 5.**
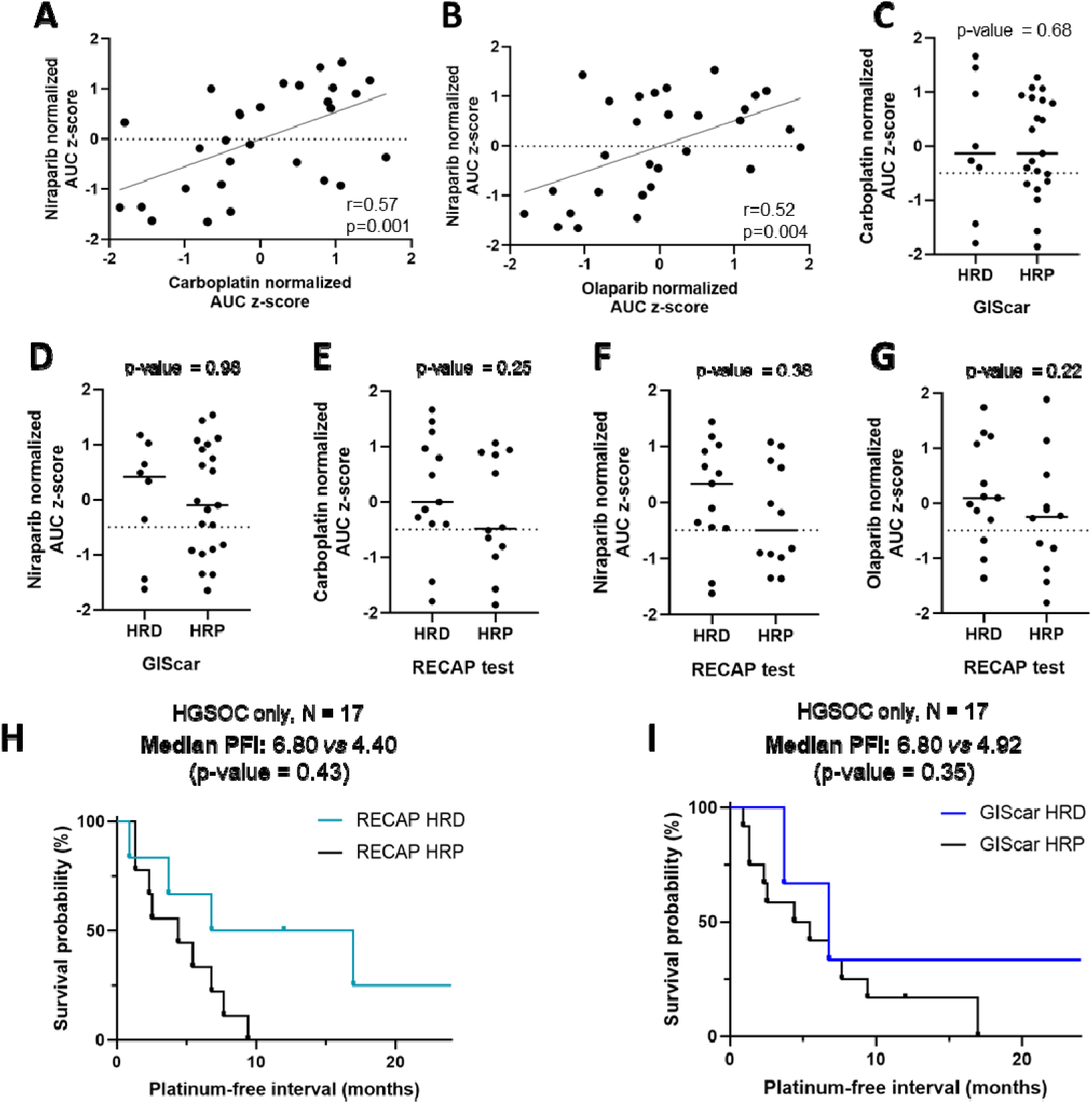
PDTO can be used to assess homologous recombination deficiency. A. Scatter plot of carboplatin and niraparib normalized AUC z-scores with the Spearman’s correlation (n=29). B. Scatter plot of olaparib and niraparib normalized AUC z-scores with the Spearman’s correlation (n=29). C, D. Dot plots comparing PDTO response to C. carboplatin or D. niraparib, expressed as normalized AUC z-score, and the genomic instability score HR status (n=29). Data were analyzed using unpaired two-sided Mann– Whitney test. E, F, G. Dot plots comparing PDTO response to E. carboplatin, F. niraparib, or G. olaparib, expressed as normalized AUC z-score, and the RECAP test HR status (n=23). Data were analyzed using unpaired two-sided Mann–Whitney test. H. Kaplan-Meier plot comparing PFI of the RECAP HRD group (n=6) and the RECAP HRP group (n=11). I. Kaplan-Meier plot comparing the PFI of the GIScar HRD group (n=4) and the GIScar HRP group (n=13).

**Supplementary Table 1.** PDTO histotypes diversity and short- and long-term establishment rates.

| Histotypes | RECEIVED AT THE LABORATORY |  | SHORT TERM ESTABLISHMENT (>p3) |  |  |  | LONG TERM ESTABLISHMENT (>p8) |  |  |  |
| --- | --- | --- | --- | --- | --- | --- | --- | --- | --- | --- |
|  |  |  | ESTABLISHED |  | % OF ESTABLISHMENT |  | ESTABLISHED |  | % OF ESTABLISHMENT |  |
|  | Number of samples | Number of patients represented | Number of samples | Number of patients represented | % of samples | % of patients represented | Number of samples | Number of patients represented | % of samples | % of patients represented |
| AGC | 15 | 9 | 1 | 1 | 6,7% | 11,1% | 0 | 0 | 0,0% | 0,0% |
| CCC | 16 | 13 | 5 | 5 | 31,3% | 38,5% | 4 | 4 | 25,0% | 30,8% |
| CS | 14 | 10 | 4 | 5 | 28,6% | 50,0% | 3 | 2 | 21,4% | 20,0% |
| EM | 18 | 10 | 5 | 4 | 27,8% | 40,0% | 1 | 1 | 5,6% | 10,0% |
| HGSOC | 261 | 170 | 50 | 40 | 19,2% | 23,5% | 26 | 23 | 10,0% | 13,5% |
| HSCC | 1 | 1 | 0 | 0 | 0,0% | 0,0% | 0 | 0 | 0,0% | 0,0% |
| LGSOC | 20 | 9 | 7 | 5 | 35,0% | 55,6% | 0 | 0 | 0,0% | 0,0% |
| LSCC | 1 | 1 | 0 | 0 | 0,0% | 0,0% | 0 | 0 | 0,0% | 0,0% |
| MC | 3 | 1 | 3 | 1 | 100,0% | 100,0% | 3 | 1 | 100,0% | 100,0% |
| TOTAL | 349 | 224 | 75 | 61 | 21,5% | 27,2% | 37 | 31 | 10,6% | 13,8% |

AGC: adult granulosa cell tumor
CCC: clear cell carcinoma
CS: carcinosarcoma
EM: endometroid carcinoma
HGSOC: High-grade serous ovarian carcinoma
HSCC: hypercalcemic small cell carcinoma
LGSOC: low-grade serous ovarian carcinoma
LSCC: lung-type small cell carcinoma
MC: mucinous carcinoma

**Supplementary Table 2.**
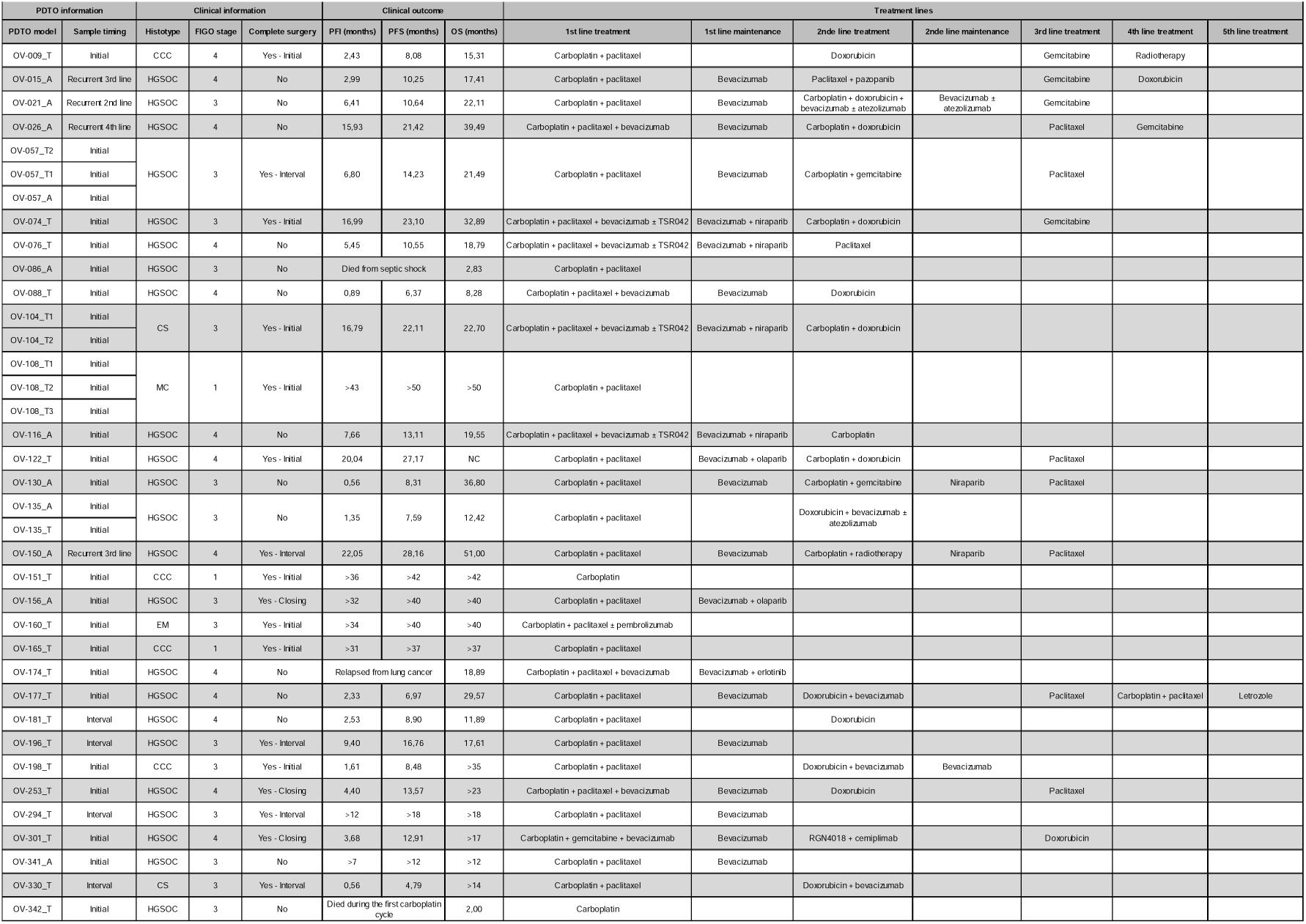
Patient clinical information. CCC: clear cell carcinoma, CS: carcinosarcoma, EM: endometroid carcinoma, HGSOC: high-grade serous ovarian carcinoma, MC: mucinous carcinoma.

**Supplementary Table 3.**
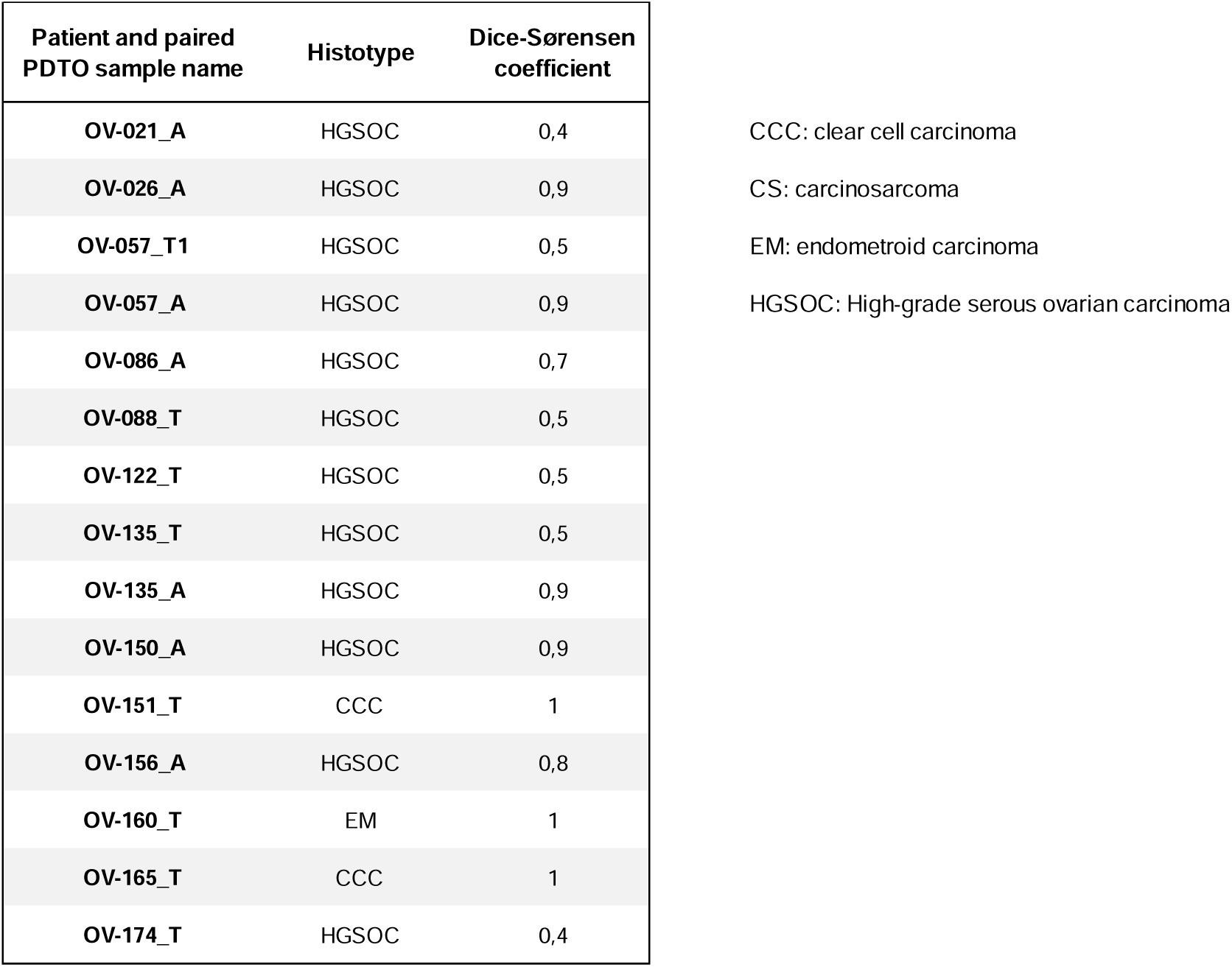
Copy-number variation (CNV) homology between paired samples.

## References

1. Cancer Today. https://gco.iarc.who.int/today/.

2. Lheureux, S., Braunstein, M. & Oza, A. M. Epithelial ovarian cancer: Evolution of management in the era of precision medicine. CA Cancer J Clin 69, 280–304 (2019).

3. Berek, J. S., Renz, M., Kehoe, S., Kumar, L. & Friedlander, M. Cancer of the ovary, fallopian tube, and peritoneum: 2021 update. Int J Gynaecol Obstet 155, 61–85 (2021).

4. González-Martín, A. et al. Newly diagnosed and relapsed epithelial ovarian cancer: ESMO Clinical Practice Guideline for diagnosis, treatment and follow-up. Ann Oncol 34, 833–848 (2023).

5. Siegel, R. L., Giaquinto, A. N. & Jemal, A. Cancer statistics, 2024. CA Cancer J Clin 74, 12–49 (2024).

6. Gulia, S. et al. Maintenance therapy with a poly(ADP-ribose) polymerase inhibitor in patients with newly diagnosed advanced epithelial ovarian cancer: individual patient data and trial-level meta-analysis. ESMO Open 7, 100558 (2022).

7. Mirza, M. R. et al. Niraparib Maintenance Therapy in Platinum-Sensitive, Recurrent Ovarian Cancer. N Engl J Med 375, 2154–2164 (2016).

8. Leman, R. et al. Validation of the Clinical Use of GIScar, an Academic-developed Genomic Instability Score Predicting Sensitivity to Maintenance Olaparib for Ovarian Cancer. Clin Cancer Res 29, 4419–4429 (2023).

9. Federici, G. & Soddu, S. Variants of uncertain significance in the era of high-throughput genome sequencing: a lesson from breast and ovary cancers. J Exp Clin Cancer Res 39, 46 (2020).

10. Fieuws, C. et al. Identification of potentially actionable genetic variants in epithelial ovarian cancer: a retrospective cohort study. NPJ Precis Oncol 8, 71 (2024).

11. Suehnholz, S. P. et al. Quantifying the Expanding Landscape of Clinical Actionability for Patients with Cancer. Cancer Discov 14, 49–65 (2024).

12. Letai, A., Bhola, P. & Welm, A. L. Functional precision oncology: Testing tumors with drugs to identify vulnerabilities and novel combinations. Cancer Cell 40, 26–35 (2022).

13. Aboulkheyr Es, H., Montazeri, L., Aref, A. R., Vosough, M. & Baharvand, H. Personalized Cancer Medicine: An Organoid Approach. Trends Biotechnol. 36, 358–371 (2018).

14. Thorel, L. et al. Patient-derived tumor organoids: a new avenue for preclinical research and precision medicine in oncology. Exp Mol Med (2024) doi:10.1038/s12276-024-01272-5.

15. Veninga, V. & Voest, E. E. Tumor organoids: Opportunities and challenges to guide precision medicine. Cancer Cell 39, 1190–1201 (2021).

16. Gorski, J. W. et al. Utilizing Patient-Derived Epithelial Ovarian Cancer Tumor Organoids to Predict Carboplatin Resistance. Biomedicines 9, 1021 (2021).

17. Hoffmann, K. et al. Stable expansion of high-grade serous ovarian cancer organoids requires a low-Wnt environment. EMBO J 39, e104013 (2020).

18. Chen, H. et al. Short-term organoid culture for drug sensitivity testing of high-grade serous carcinoma. Gynecologic Oncology (2020) doi:10.1016/j.ygyno.2020.03.026.

19. Hill, S. J. et al. Prediction of DNA Repair Inhibitor Response in Short-Term Patient-Derived Ovarian Cancer Organoids. Cancer Discov 8, 1404–1421 (2018).

20. de Witte, C. J. et al. Patient-Derived Ovarian Cancer Organoids Mimic Clinical Response and Exhibit Heterogeneous Inter- and Intrapatient Drug Responses. Cell Reports 31, 107762 (2020).

21. Maru, Y., Tanaka, N., Itami, M. & Hippo, Y. Efficient use of patient-derived organoids as a preclinical model for gynecologic tumors. Gynecol Oncol 154, 189–198 (2019).

22. Kopper, O. et al. An organoid platform for ovarian cancer captures intra- and interpatient heterogeneity. Nat Med 25, 838–849 (2019).

23. Phan, N. et al. A simple high-throughput approach identifies actionable drug sensitivities in patient-derived tumor organoids. Commun Biol 2, 78 (2019).

24. Senkowski, W. et al. A platform for efficient establishment and drug-response profiling of high-grade serous ovarian cancer organoids. Dev Cell 58, 1106–1121.e7 (2023).

25. Hu, H., Sun, C., Chen, J. & Li, Z. Organoids in ovarian cancer: a platform for disease modeling, precision medicine, and drug assessment. J Cancer Res Clin Oncol 150, 146 (2024).

26. DeLair, D. et al. HNF-1β in ovarian carcinomas with serous and clear cell change. Int J Gynecol Pathol 32, 541–546 (2013).

27. Naipal, K. A. T. et al. Functional ex vivo assay to select homologous recombination-deficient breast tumors for PARP inhibitor treatment. Clin. Cancer Res. 20, 4816–4826 (2014).

28. Maenhoudt, N. et al. Developing Organoids from Ovarian Cancer as Experimental and Preclinical Models. Stem Cell Reports 14, 717–729 (2020).

29. Nanki, Y. et al. Patient-derived ovarian cancer organoids capture the genomic profiles of primary tumours applicable for drug sensitivity and resistance testing. Sci Rep 10, 12581 (2020).

30. Bi, J. et al. Successful Patient-Derived Organoid Culture of Gynecologic Cancers for Disease Modeling and Drug Sensitivity Testing. Cancers (Basel) 13, 2901 (2021).

31. Ford, C. E., Werner, B., Hacker, N. F. & Warton, K. The untapped potential of ascites in ovarian cancer research and treatment. Br J Cancer 123, 9–16 (2020).

32. Arias-Diaz, A. E. et al. Ascites-Derived Organoids to Depict Platinum Resistance in Gynaecological Serous Carcinomas. Int J Mol Sci 24, 13208 (2023).

33. Ray-Coquard, I. et al. Olaparib plus Bevacizumab as First-Line Maintenance in Ovarian Cancer. N Engl J Med 381, 2416–2428 (2019).

34. Thorel, L. et al. The OVAREX study: Establishment of ex vivo ovarian cancer models to validate innovative therapies and to identify predictive biomarkers. BMC Cancer 24, 701 (2024).

35. Talevich, E., Shain, A. H., Botton, T. & Bastian, B. C. CNVkit: Genome-Wide Copy Number Detection and Visualization from Targeted DNA Sequencing. PLOS Computational Biology 12, e1004873 (2016).

36. Bankhead, P. et al. QuPath: Open source software for digital pathology image analysis. Sci Rep 7, 16878 (2017).

37. Stringer, C., Wang, T., Michaelos, M. & Pachitariu, M. Cellpose: a generalist algorithm for cellular segmentation. Nat Methods 18, 100–106 (2021).

38. Schmidt, U., Weigert, M., Broaddus, C. & Myers, G. Cell Detection with Star-Convex Polygons. in Medical Image Computing and Computer Assisted Intervention – MICCAI 2018 (eds. Frangi, A. F., Schnabel, J. A., Davatzikos, C., Alberola-López, C. & Fichtinger, G.) 265–273 (Springer International Publishing, Cham, 2018). doi:10.1007/978-3-030-00934-2_30.

39. van der Walt, S. et al. scikit-image: image processing in Python. PeerJ 2, e453 (2014).

40. scikit-image’s documentation — skimage 0.25.0rc2.dev0 documentation. https://scikit-image.org/docs/dev/index.html.

41. Xuan, G., Zhang, W. & Chai, P. EM algorithms of Gaussian mixture model and hidden Markov model. in Proceedings 2001 International Conference on Image Processing (Cat. No.01CH37205) vol. 1 145–148 vol.1 (2001).

42. Jolliffe, I. T. Principal Component Analysis and Factor Analysis. in Principal Component Analysis (ed. Jolliffe, I. T.) 115–128 (Springer, New York, NY, 1986). doi:10.1007/978-1-4757-1904-8_7.

